# Recurring *EPHB1* mutations in human cancers alter receptor signalling and compartmentalisation of colorectal cancer cells

**DOI:** 10.1101/2023.03.18.533270

**Authors:** Snehangshu Kundu, Luís Nunes, Jeremy Adler, Lucy Mathot, Ivaylo Stoimenov, Tobias Sjöblom

**Author notes:** To whom correspondence should be addressed, at. These authors contributed equally (Snehangshu Kundu, Luís Nunes).

## Abstract

Ephrin (EPH) receptors have been implicated in tumorigenesis and metastasis, but the functional understanding of mutations observed in human cancers is limited. We previously demonstrated reduced cell compartmentalisation for somatic *EPHB1* mutations found in metastatic colorectal cancer cases. We, therefore, integrated pan-cancer and pan-EPH mutational data to prioritise recurrent *EPHB1* mutations for functional studies to understand their contribution to cancer development and metastasis. Here, 79,151 somatic mutations in 9,898 samples of 33 different tumour types were analysed to find 3D-mutated cluster pairs and recurring hotspot mutations in EPH receptors. From these, 15 recurring *EPHB1* mutations were stably expressed in colorectal cancer cells. Whereas the ligand-binding domain mutations C61Y, R90C, and R170W, the fibronectin domain mutation R351L, and the kinase domain mutation D762N displayed reduced to strongly compromised cell compartmentalisation, the kinase domain mutations R743W and G821R enhanced this phenotype. While mutants with reduced compartmentalisation also had reduced ligand induced receptor phosphorylation, the enhanced compartmentalisation was not linked to receptor phosphorylation level. Phosphoproteome mapping pinpointed the PI3K pathway and PIK3C2B phosphorylation in cells harbouring mutants with reduced compartmentalisation. This is the first integrative study of pan-cancer EPH receptor mutations followed by *in vitro* validation, a robust way to identify cancer-causing mutations.

## Introduction

Receptor tyrosine kinases (RTKs) play important roles in cell proliferation, differentiation, and motility (1,2). The ephrin (EPH) receptor is the largest subfamily of RTKs with 14 members classified into subtype A (*EPHA1*-*8* and *EPHA10*) and B (*EPHB1*-*4* and *EPHB6*). Human EPHA receptors preferentially bind ephrin (EFN) A1-A5 ligands, whereas the EPHB receptors bind EFNB1-B3 ligands, respectively. EPH-EFN signalling is initiated by polymerisation of EFN ligand bound EPH receptors and is eventually attenuated by condensation through coalescence of many polymerized clusters (3). In contrast to the unidirectional signalling of other RTKs, the binding of EFN ligand to EPH receptors can also initiate bidirectional communication with forward signalling in the receptor-expressing cells and retrograde signalling in the ligand-expressing cells (4). Deregulation of EPH expression has been linked to both pro- and anti-tumorigenic properties in different tumour types (2). Somatic mutations in cancer have been shown to modulate EPH function (5-7). Thus, identification and functional characterization of somatic EPH mutations can advance the understanding of their roles in tumour development.

Approximately 20-25% of colorectal cancer (CRC) patients present with metastatic disease at diagnosis, and another 20-25% will end up developing metastasis later in the course of the disease. The lack of efficient treatments for metastatic disease leads to a high overall 40-45% mortality rate (8,9). Identifying patients that require close monitoring to detect recurrence as well as to stratify patients that would benefit most from adjuvant chemotherapy treatment is of clinical importance. Despite the recent progress in cancer genome sequencing, it has proven challenging to associate specific gene mutations with metastasis. For example, *FBXW7* has been proposed to be preferentially mutated in non-metastatic cases (10,11), whereas loss of 1p36 has been associated with metastasis of CRC (12). The EPH receptors have been linked to metastatic disease due to their roles in tumour growth, invasiveness, angiogenesis, and metastasis. Reduced *EPHB1* expression in colon cancer was associated with poor differentiation and increased invasive capacity (13). We recently demonstrated a link between *EPHB1* inactivating mutations and metastasis of primary CRC (5). Two *EPHB1* mutations exhibited compromised cell repulsion with *EFNB1* ligand-expressing cells in an *in vitro* compartmentalisation assay (14) as compared to wild-type *EPHB1* (5). This warranted further studies of the contribution of EPH receptor mutations to CRC development and metastasis. Due to the intermediate-low mutation frequencies of EPH receptors in cancer, we integrated mutational data from several tumour types and different EPH receptors to build evidence for recurrent mutated positions worthy of further functional studies, evaluated selected hotspot mutations based on the compartmentalisation phenotype and determined the impact of mutations on EPH receptor phosphorylation.

## Material and Methods

### Identification and prioritisation of tumour-derived EPH receptor mutants for functional studies

To select the most relevant *EPHB1* mutations, somatic mutation data for EPH receptors was obtained from The Cancer Genome Atlas (TCGA) portal for 33 different tumour types (https://www.cancer.gov/tcga; data accessed in January 2017). From the putative somatic mutations retrieved, we retained those which had (i) a tumour and matching normal sequence coverage of more than 30 reads, (ii) more than 10% of alleles in the tumour sample supporting the variant sequence and (iii) more than 99% of alleles in the normal sample supporting the reference sequence. The mutations were mapped to the Consensus Coding Sequence (CCDS, release 20) transcripts definition with GRCh38 used as the reference human genome (15). Canonical protein sequences retrieved from the UniProt database (16) were used to transform exon-coding variant genomic coordinates into protein coordinates. Coding non-synonymous mutations were then annotated for functional impact using ANNOVAR dbNSFP version 3.3a (17). A customised score was given to each amino acid alteration, by averaging the outputs of ten different functional impact prediction algorithms (Supplementary Table 1). All canonical EPH receptor sequences were aligned with UniProt alignment tools to produce a single consensus sequence. From the resulting alignment, the sum of the different EPH receptor mutations was calculated for each aligned position in the consensus sequence. HotSpot3D-1.3.11 (18) was used to analyse mutational hotspots in the 3D-structure of the EPH receptors, from crystal X-ray diffraction and solution nuclear magnetic resonance structures (Supplementary Table 2). The algorithm was used with default parameters to output amino acid pair proximities based on the average distance between the residues. Amino acids pairs were selected if they (i) had statistically significant interaction (*p* ≤ 0.05), (ii) were in the same protein chain and (iii) were separated by at least five amino acids in the linear protein sequence. Each cluster pair position was then transformed into canonical EPH receptor positions by mapping to the consensus sequence. We next considered mutations with average functional impact score ≥ 6. A 3D-mutation cluster pair was selected if (i) the 3D-mutation partner had more than one mutation, (ii) the pair was > 5 Å apart in the structure and (iii) > 50 amino acids apart in the linear protein sequence (Figure 1A). Mutations without a mutation partner in the 3D-protein structure were sorted by prevalence and prioritised by predicted average functional impact. Finally, cluster pairs and mutated positions were selected if they had at least one *EPHB1* mutation. If the selected mutation had more than one amino acid change, amino acid physical properties and functional impact were assessed for each alteration and the one with the higher predicted impact was selected.

**Figure 1.**
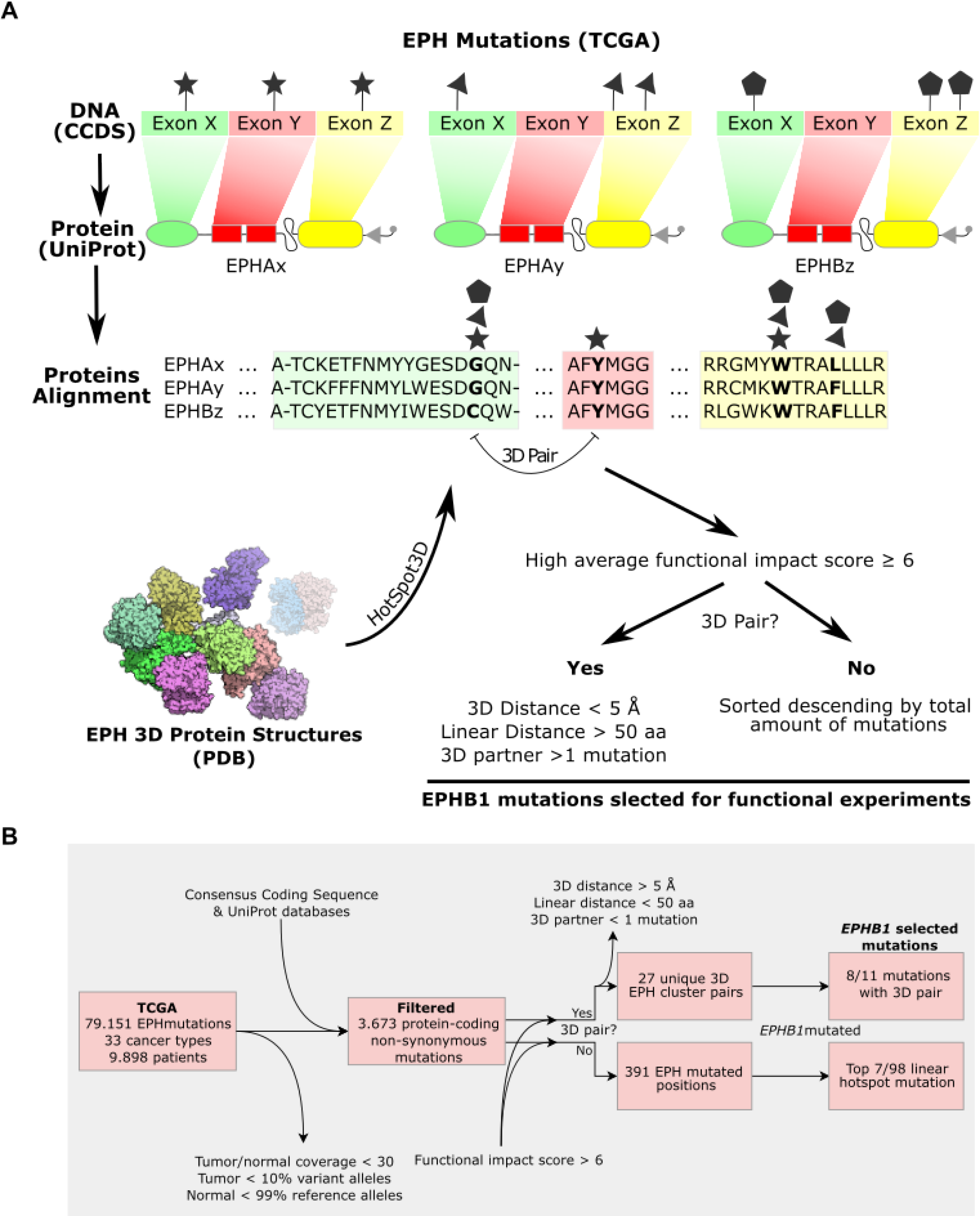
Selection of recurring *EPHB1* mutations from pan-ephrin TCGA data from 9,989 patients in 33 different cancer types. A. Mutations were filtered and mapped onto each respective EPH coding sequence from The Consensus Coding Sequence (CCDS) database. DNA positions were transformed into each canonical EPH protein sequence from the UniProt database and aligned to project all mutations onto a consensus EPH receptor sequence. Mutation pairs proximal in 3D but distant in the linear protein sequence were identified using HotSpot3D employing all available EPH 3D-structures reported in The Protein Databank (PDB) database. Non-synonymous mutated positions in *EPHB1* with high functional impact were selected based on distance if they were in a 3D-pair cluster or if not by prevalence of mutations. B. Flow chart of the *EPHB1* mutation selection.

### Cell lines and cell culture

Parental DLD-1 (CCL-221) colorectal cells were purchased from ATCC (USA) and authenticated by short tandem repeat profiling using the ATCC cell line authentication service. All cell cultures were maintained in McCoy’s 5A medium (Thermo Fisher Scientific, USA) supplemented with 10% fetal bovine serum and 1% penicillin-streptomycin (Thermo Fisher Scientific, USA) at 37ºC in 5% CO_2._

### Lentiviral constructs

Lentiviruses expressing custom-designed vector backbone pReceiver-Lv225 with *EPHB1* wildtype or mutants plus eGFP marker, and pReceiver-Lv224 with *EFNB1* wildtype plus mCherry marker were purchased from LabOmics (Belgium). Both fluorescent markers, eGFP and mCherry, were not tagged directly with either *EPHB1* receptor or *EFNB1* ligand as both the markers were present downstream of IRES2 (Supplementary Figure 1). Hence, both fluorescent marker proteins were expressed individually. Negative control constructs expressing only eGFP and mCherry were also acquired.

### Generation of stable cell lines

Lentivirus transduction of DLD-1 CRC cells was used to achieve stable overexpression of EFNB1 ligand, wild-type EPHB1 or mutant EPHB1. The day before transduction, 50,000 cells were plated in each well of a 24-well plate. Viruses were diluted in 250 µl of normal growth medium with 7.5 μg/ml Sequa-Brene (Merck KGaA, Germany) per well. The plating medium was removed and 250 µl of diluted virus was added to each well. After 24 h incubation at 37ºC, virus-containing media were replaced with fresh medium. After 48 h incubation, transduced cells were selected with puromycin (1 µg/ml) for 2-4 passages to remove puromycin non-resistant cells from the cell pool.

### Detection and identification of EPHB1 mutant transcripts by Sanger sequencing

Total RNA was isolated from DLD-1 cells ectopically expressing *EPHB1* mutants using the RNeasy Plus Mini Kit (Qiagen, Germany). First strand cDNA synthesis was conducted with the RevertAid H-minus First Strand cDNA synthesis kit (Thermo Fisher Scientific, USA). Next, PCR amplification was performed with cDNA as template using customised sequencing primers (Supplementary Table 3). PCR was performed in 20 µl reactions containing 1 × Phusion HF buffer (Thermo Fisher Scientific, USA); 0.2 mM dNTPs (Thermo Fisher Scientific, USA), 0.5 µM forward and reverse primers (Merck KGaA, Germany), 0.02 U Phusion Hot Start II High-Fidelity DNA polymerase (Thermo Fisher Scientific, USA) and 10-50 ng cDNA. Reactions were carried out in a thermocycler using the following PCR protocol: 98ºC for 30 s; 30 cycles: 98ºC for 10 s, 64º for 15 s, and 72ºC for 20 s; 72ºC for 10 min. PCR products were sent for Sanger sequencing with the customised primers at Eurofins Genomics Europe (Germany).

### Detection of EPHB1 mutant proteins

Pellets from 2×10^6^ cells were lysed in RIPA buffer (Thermo Fisher Scientific, USA) and protein concentration was estimated by the Pierce BCA Protein assay kit (Thermo Fisher Scientific, USA). For protein separation, 15 µg of each sample was loaded on a NuPage 4-12% Bis-Tris gel (Thermo Fisher Scientific, USA) and run at 180 V for 1 h. Proteins were transferred to a membrane using nitrocellulose iBlot transfer stacks (Thermo Fisher Scientific, USA) at 20V for 8 min. The membrane was cut into three parts according to the size of the respective proteins to be detected. The top part of the membrane was blocked with 3% non-fat milk in Tris-buffered saline with 0.1% Tween (TBST) followed by incubation at room temperature (RT) for 1 h with mouse monoclonal anti-FLAG primary antibody (Sigma-Aldrich, USA; #F3165, 1:2,000) to detect the overexpressed *EPHB1* receptors proteins (130 KDa). The middle part of the membrane was blocked with 5% bovine serum albumin (BSA) in TBST followed by incubation with mouse monoclonal anti-β-Actin (Santa Cruz Biotechnology, USA; #47778, 1:10,000) primary antibody at RT for 1 h as loading control. The lower part of the membrane was blocked with 3% BSA in phosphate-buffered saline (PBS) solution followed by incubation with rabbit polyclonal anti-GFP (Abcam, UK; #ab290, 1:1,000) primary antibody at RT for 1 h for eGFP detection (30 kDa). After incubation with primary antibodies, the top and middle membranes were incubated at RT for 1 h with HRP-conjugated anti-mouse secondary antibody (Thermo Fisher Scientific, USA; #31430, 1:8,000) and the lower part of the membrane with HRP-conjugated anti-rabbit secondary antibody (Cytiva, USA; #NA934, 1:6,000).

### Cell compartmentalisation-based-phenotypic screen

To identify *EPHB1* mutations with compartmentalisation phenotypes (5), *in vitro* co-cultures of eGFP labelled cells expressing no receptor, wild-type or mutant *EPHB1* with mCherry labelled cells expressing either no ligand or *EFNB1* were plated in four replicates in 96-well plates at a receptor to ligand-expressing cell ratio of 1:3. The experiment was imaged in real-time using an IncuCyte2016A (Essen BioScience, USA) for 120-180 h. Background due to green autofluorescence was subtracted by applying Top-Hat on green clusters. Subsequently, a custom-designed processing definition was applied to analyse the data. The screens were repeated at least 3 times.

### Confocal microscopy-based in vitro compartmentalisation assays

Confocal microscopy-based compartmentalisation experiments were performed as described previously (5,14). Briefly, DLD-1 cells expressing different *EPHB1* wildtype or mutant versions and *EFNB1* were mixed in suspension at a ratio of 1:3 and plated at a density of 130,000 cells/cm^2^ on coverslips coated with 2 mg/cm^2^ of 1-2 mg/mL laminin (Merck KGaA, Germany, USA) and incubated at 37ºC in 5% CO_2_. The respective negative controls were eGFP or mCherry expressing cells. Culture medium was changed after 24 h, and after 48 h, the coverslips were fixed in 4% paraformaldehyde for 20 min at RT and mounted in DAPI Fluoromount-G (SouthernBiotech, USA). Slides were subjected to confocal imaging in a Zeiss LSM 700 microscope (Zeiss, Germany) and compartmentalisation was quantified by counting the percentage of total green cells in each eGFP-positive cluster of 10 representative fields under 20× NA 0.8 objective at two different confocal planes in the z-axis and from two experimental repeats using a custom ImageJ macro. Images throughout the paper show the basal plane. The experiments were performed twice. Statistical analyses were performed using the Mann-Whitney *U* test.

### Ephrin receptor phosphorylation following stimulation with ephrin B1 ligand

The EPHB1 receptor stimulation was performed using pre-clustered EphrinB1-Fc fragment (Cortina et al., 2007). Briefly, 5×10^5^ cells were plated at 50% confluency in a 6-well plate and incubated overnight. The next day, cells were washed once with HBSS and starvation medium (0.1% BSA) was added and incubated for 24 h at 37ºC in 5% CO_2_. For cluster formation, EphrinB1-Fc (R&D systems, USA) and control Fc fragment (R&D Systems; USA) were mixed with purified anti-Fc-antibody in 2:1 molar ratio and incubated at RT for 2 h. After starvation, 1 ml of fresh starvation medium with 0.5 µM of pre-clustered EphrinB1-Fc and Fc fragment was added to each well followed by incubation at 37°C for 30 min. After the stimulation, the plate was immediately transferred on ice, washed once with ice-cold HBSS and the cells lysed in 600 µl of RIPA lysis buffer (Thermo Fisher Scientific, USA) to prepare the cell lysates for immunoblot analysis. Phosphorylated EPHB1 was detected with anti-phospho-T594/604-EPHB1-ab (Merck KGaA, Germany; USA; #SAB4504172, 1:5,000) and total EPHB1 with anti-FLAG primary antibody (Merck KGaA, Germany; USA; #F3165, 1:20,000) with SuperSignal Western Blot Enhancer (Thermo Fisher Scientific, USA) and quantitated using Image J. Each experiment was repeated at least 2 times.

### Proteome and phospho-proteome analysis

Wild-type EPHB1 expressing DLD-1 cells along with four mutants (C61Y, D762N, R351L and R743W) were subjected to analysis. Cells were stimulated with 0.5 µM of pre-clustered EphrinB1-Fc ligand for 30 min, followed by protein extraction in scioExtract buffer (Sciomics, Germany). Each condition had three technical replicates. After quality control of the samples, bulk protein concentration was determined by BCA protein assay. The samples were labelled at an adjusted protein concentration for 2 h with scioDye 2 (Sciomics, Germany), followed by removal of excess dye and buffer exchange to PBS. The labelled protein samples were stored at -20°C until analysis on 18 scioDiscover antibody microarrays (Sciomics, Germany). Each antibody was represented in four replicates on the arrays. The arrays were blocked with scioBlock (Sciomics, Germany) on a Hybstation 4800 (Tecan, Austria) followed by incubation with scioPhosphomix 1 (Sciomics, Germany). After incubation, the slides were thoroughly washed with 1 × PBSTT, rinsed in 0.1×PBS and ddH_2_O and subsequently dried with nitrogen. Slide scanning was conducted using a Powerscanner (Tecan, Austria) with constant instrument laser power and PMT settings. Spot segmentation was performed with GenePix Pro 6.0 (Molecular Devices, USA). Acquired raw data were analysed using the linear models for microarray data (LIMMA) package of R-Bioconductor after uploading the median signal intensities. For normalisation, Cyclic Loess normalisation was applied. For analysis of the samples, a one-factorial linear model was fitted via least squares regression with LIMMA, resulting in a two-sided t-test or F-test based on moderated statistics. All presented *p* values were adjusted for multiple testing by controlling the false discovery rate according to Benjamini and Hochberg. Differences in protein abundance or phosphorylation level between different samples or sample groups are presented as log-fold changes (logFC) calculated for the basis 2.

## Results

### A compendium of somatic mutations in EFN ligands and EPH receptors in human cancers

To discover mutational hotspots in the highly homologous EPH receptors, we analysed 79,151 putative somatic mutations from 33 different tumour types in a total of 9,898 patients (Supplementary Table 4). After removal of low confidence variants and non-coding and synonymous mutations, 3,673 protein-coding non-synonymous variants remained (5% of total variants). Of these, 3,009 were missense, 2 stop-loss, 244 nonsense, 73 frameshift insertions, 166 frameshift deletions, 35 in-frame indels, 142 splice-site and 2 unknown variants. For each EPH receptor subtype, coding non-synonymous mutations were more common in *EPHA3* and *EPHB1*, with 0.4 and 0.33 mutations per amino acid, respectively. Cutaneous melanoma, diffuse large B-cell lymphoma and lung adenocarcinoma had EPH receptor alteration frequencies of 51.1%, 40.4% and 38.4% of cases, respectively. When considering only *EPHB1*, the most frequent altered tumour types included uterine corpus endometrial carcinoma, lung, and colon adenocarcinoma with mutation frequencies of 10.9%, 6.3% and 6%, respectively (Supplementary Table 5).

First, we sought to identify hotspots in 3D-space that might be overlooked in conventional analyses based on mutation density in the linear sequence. From the predicted high functional impact mutations, after filtering proximal amino acid pairs by distance in protein sequence and structure, we found 27 unique EPH 3D-cluster pairs (Figure 1B). When considering only those with mutations in *EPHB1*, 11 cluster pairs remained (Supplementary Table 6). Of these, 3 positions were excluded from further studies, L709 due to conservative amino acid change (leucine to isoleucine), and V391 and M23 as they had *EPHB1* mutations in single tumours only and were located in non-conserved positions. Of the 8 remaining positions, 4 resided in the ligand-binding domain and 4 in the kinase domain. In total, 391 EPH positions were mutated without a 3D-partner amino acid associated. When considering only positions mutated in *EPHB1*, 98 were retained and the 7 most prevalent considering all EPH receptor mutations were selected for further studies (Supplementary Table 7). The position D374 was excluded as it was a non-conserved position. Of the 7 selected mutations, 1 was located in the fibronectin type-III 1 domain, 4 in the kinase domain, and 2 outside of known domains. In total, 15 *EPHB1* mutations were selected for functional characterisation (Table 1 and Supplementary Figure 3A).

**Table 1.** *EPHB1* mutations identified in 9,898 cases of 33 different TCGA tumour types. TCGA pan-cancer pan-EPH somatic mutations were filtered and *EPHB1* coding non-synonymous mutations with a high functional impact were selected if they either had (i) another cognate mutated amino acid in close proximity in 3D-space or (ii) by mutation prevalence. Primary in vitro phenotypical screening by IncuCyte identified the mutations with compartmentalisation phenotype for validation by confocal microscopy. **(*)** Mutants selected for confocal analyses. The position **(a)** V248M is located between the ligand-binding and the fibronectin type-III 1 domains in a compositional bias region enriched for cysteine; and **(b)** R883Q is located in the position C-terminally to the protein kinase domain. n/a, mutated positions were not associated with another amino acid in the 3D space.

| *EPHBI*<br>Amino acid<br>Position | *EPHBI*<br>Mutations | Total EPH<br>Mutations | 3D-Partner<br>Position<br>(*EPHBI*) | Protein Domain |
| --- | --- | --- | --- | --- |
| G685D* | 1 | 4 | I696 | Protein Kinase |
| R743W* | 5 | 13 | F801 | Protein Kinase |
| G821R* | 2 | 7 | R170 | Protein Kinase |
| C61Y* | 1 | 2 | C183 | Ligand Binding |
| R90C* | 2 | 6 | S188 | Ligand Binding |
| R682C | 3 | 10 | E698 | Protein Kinase |
| T117I* | 2 | 3 | G172 | Ligand Binding |
| R170W* | 7 | 9 | E116 | Ligand Binding |
| V248M | 1 | 26 | n/a | (a) |
| R883Q | 1 | 18 | n/a | (b) |
| R748S* | 2 | 14 | n/a | Protein Kinase |
| D762N* | 6 | 14 | n/a | Protein Kinase |
| R865W* | 3 | 13 | n/a | Protein Kinase |
| R351L* | 2 | 11 | n/a | Fibronectin type-III 1 |
| R799H | 1 | 10 | n/a | Protein Kinase |

We next engineered DLD1 CRC cells to overexpress eGFP alone, wildtype *EPHB1*, or each of the selected mutants by lentiviral transduction with a polycistronic lentiviral expression vector in which expression of EPHB1 and eGFP were uncoupled due to the presence of IRES elements (Supplementary Figure 1). The expression of *EPHB1* mutant transcripts was confirmed by Sanger sequencing (Supplementary Figure 2) and mutant protein was determined by immunoblotting (Supplementary Figure 3B). Notably, the expression levels of G685D and C183Y mutant EPHB1 proteins were lower than those of other EPHB1 mutants. However, the level of eGFP overexpression was similar in all cell lines, suggesting a shorter half-life for these mutant EPHB1 proteins (Supplementary Figure 3B).

### Compartmentalisation phenotypes of selected EPHB1 mutants

To investigate the functional impact of the selected *EPHB1* mutations in cancer development and metastasis, we used the *in vitro* compartmentalisation assay in human DLD-1 CRC cells. In this assay, *EPHB1* receptor-bearing cells form large homogeneous cell clusters of > 50 cells upon contact with co-cultured ephrin ligand-expressing cells (5,14). To assess the mutants based on the *EPHB1-EFNB1*-mediated compartmentalisation, a primary screen was performed using the IncuCyte real-time-imaging system (see Material and Methods). Cells expressing *EPHB1* V248M, R682C, R799H and R883Q mutants showed no compartmentalisation difference when compared to *EPHB1* wild-type cells. Mutants with enhanced or compromised compartmentalisation phenotypes relative to wild-type *EPHB1* when co-cultured with *EFNB1* ligand expressing cells were subjected to confocal microscopy-based compartmentalisation assay. Here, the mutants were divided by compartmentalisation phenotype into four groups. Phenotypes similar to that of wildtype *EPHB1* were observed for T117I, R748S and R865W (Supplementary Figure 4A and B). Reduced compartmentalisation characterized *EPHB1* R90C, R170W, R351L and D762N mutants (*p* < 0.01, < 0.01, < 0.001 and = 0.0159, respectively (Figure 2A and B). Strongly compromised compartmentalisation, essentially lacking large clusters, was observed for *EPHB1* C61Y (*p* < 0.0001; Figure 3A and B). Finally, enhanced compartmentalisation was observed for *EPHB1* R743W and G821R cells, with significantly more large clusters (>50 cells) when co-cultured with *EFNB1* ligand-expressing cells (*p* ≤ 0.01; Supplementary Figure 5A and B). Taken together, 2 mutants had enhanced cell compartmentalisation, 6 had reduced, and 7 mutants were similar to wild-type EPHB1.

**Figure 2.**
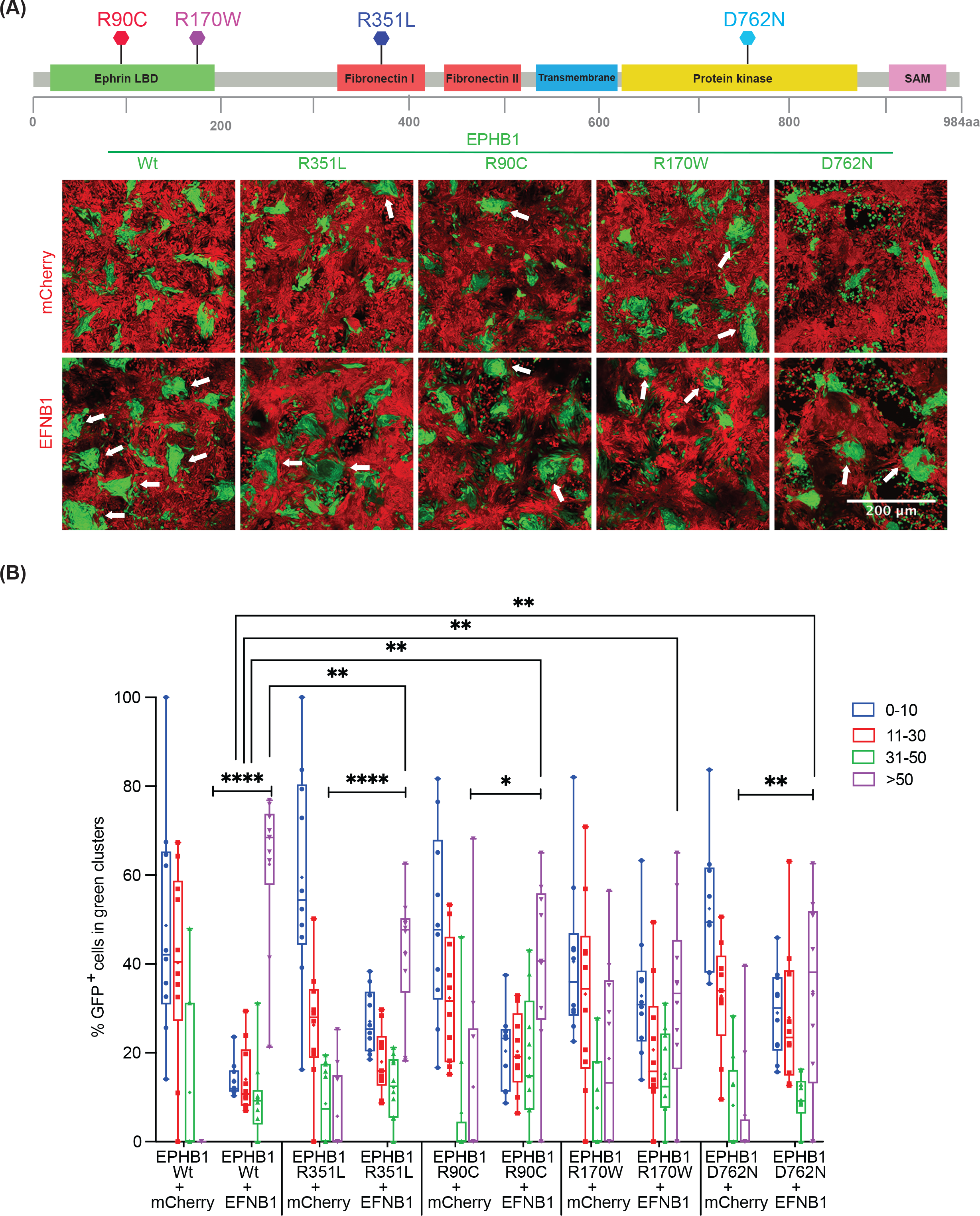
Colorectal cancer cells with *EPHB1* receptor mutations R90C, R170W, R351L and D762N had reduced compartmentalisation in presence of *EFNB1*. (A) Upper panel, positions of the mutants in the full length EPHB1 protein. Lower panel, *in vitro* compartmentalisation assays by co-culturing DLD-1 cells expressing eGFP along with either wild-type EPHB1 or R90C, R170W, R351L and D762N mutants with DLD-1 cells expressing mCherry with or without EFNB1 ligand. Representative confocal images from 10 randomly chosen image fields for each type. Arrows, examples of large (>50 cells), homogeneous GFP^+^ cell clusters indicative of cell sorting and compartmentalisation. (B) Quantitative results from the compartmentalisation experiments. Cell distribution was quantified by counting the percentage of GFP^+^ cells forming clusters of different sizes. In co-cultures of EFNB1 ligand with EPHB1 and its four mutated versions R90C, R170W, R351L and D762N, significantly lower percentage of GFP^+^ cells were distributed into large homogeneous clusters (>50) as compared to the Wt. Mann-Whitney U test was used to calculate large cluster (>50 cells) difference between each mutant with or without EFNB1 against the respective positive control condition with EPHB1 wild-type. This experiment was performed at least twice and imaged with five random fields from each experiment. Here, * *p* < 0.05, ** *p* < 0.01, *** *p* < 0.001 and **** *p* < 0.0001.

**Figure 3.**
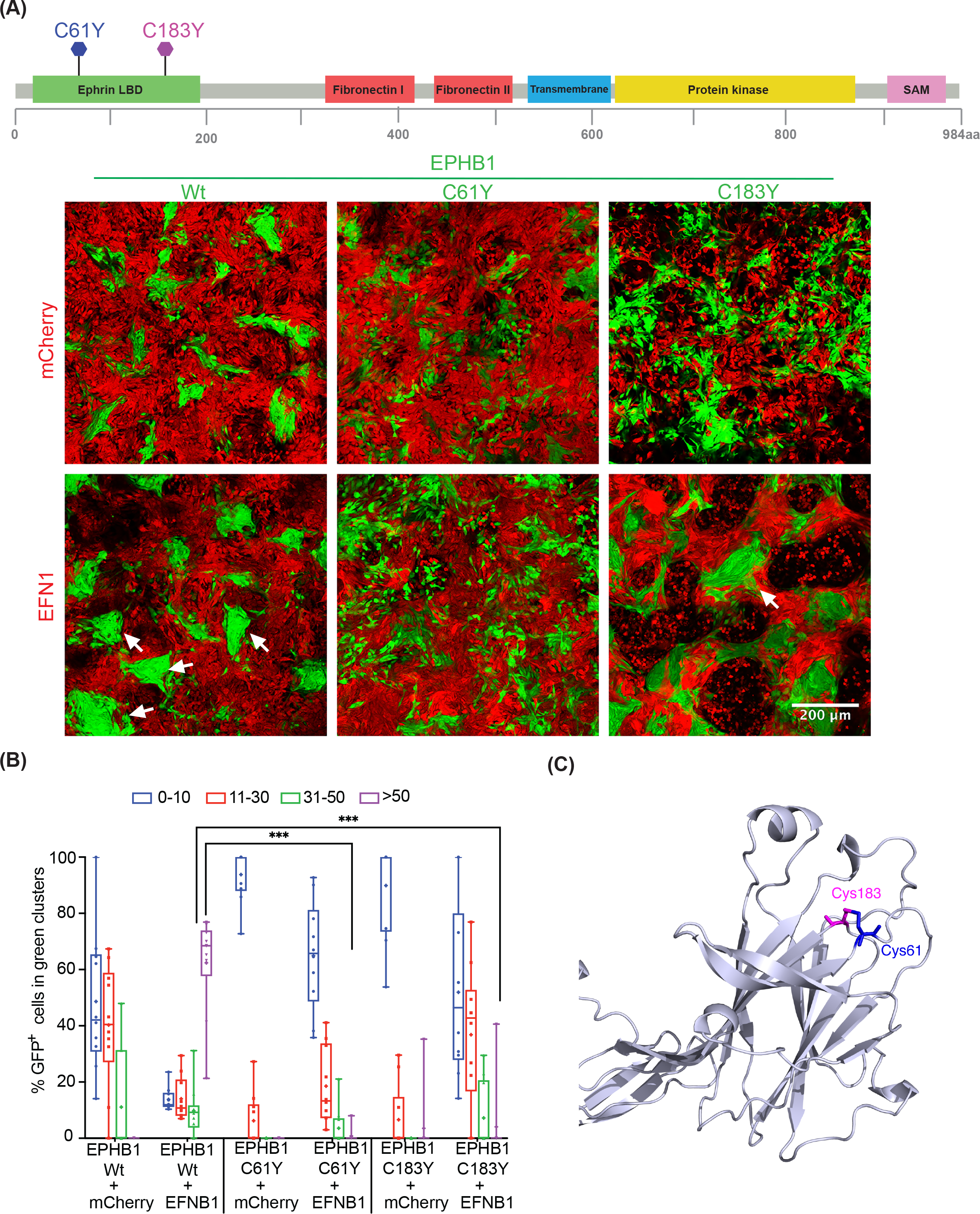
The *EPHB1* receptor 3D mutation partners C61Y and C183Y display compromised compartmentalisation in presence of *EFNB1*. (A) Upper panel, positions of the mutants on the full length EPHB1 protein. Lower panel, *in vitro* compartmentalisation assays by co-culturing DLD-1 cells expressing eGFP along with either wild-type EPHB1 or C61Y or C183Y mutants with DLD-1 cells expressing mCherry with or without EFNB1 ligand. Representative confocal images from 10 randomly chosen image fields for each type. Arrows, examples of large (>50 cells), homogeneous GFP^+^ cell clusters indicative of cell sorting and compartmentalisation. (B) Quantitative results from the compartmentalisation experiments. Cell distribution was quantified by counting the percentage of GFP^+^ cells forming clusters of different sizes. In co-cultures of EFNB1 ligand with EPHB1 and its two mutated versions (C61Yand C183Y), almost none of the GFP^+^ cells were distributed into large homogeneous clusters (>50). The Mann-Whitney *U* test was used to calculate large cluster (>50 cells) difference between each mutant with or without EFNB1 against the respective positive control condition with EPHB1 wild-type. This experiment was performed at least twice and imaged with five random fields from each experiment. Here, * *p* < 0.05, ** *p* < 0.01, *** *p* < 0.001 and **** *p* < 0.0001. (C) The positions of C61Y and C183Y in the structure of the <u>E</u>phrin <u>B</u>inding <u>D</u>omain (EBD) of EPHB1 from the Alpha-fold database.

### Ligand induced phosphorylation of EPHB1 mutant receptors

Phosphorylation of cytoplasmic tyrosine residues is a hallmark of Eph receptor activation and its downstream signal transduction (19). The phosphorylation of conserved tyrosine residues 594 and 604 of EPHB1 has been implicated in activation of ERK signalling (20). For T117I, R748S and R865W mutants, which had similar compartmentalisation phenotype as wild-type EPHB1, the ratio of phosphorylated to total EPHB1 after ligand stimulation was similar to that of wild-type EPHB1 (Figure 4A). In addition, receptor phosphorylation level after ligand stimulation was abolished for C61Y (Figure 4B) having a compromised compartmentalisation phenotype (Figure 3A-B). Significant reduction of phosphorylation after ligand stimulation (Figure 4C) was also observed for R90C, R170W, D762N and R351L EPHB1 mutants with reduced compartmentalisation (Figure 2A-B). Intriguingly, the R743W and G821R mutants with enhanced compartmentalisation (Supplementary Figure 5A and B) showed no or reduced receptor phosphorylation as compared to wild-type EPHB1 (Supplementary Figure 5C and D) after ligand stimulation, suggesting that this phenotype is independent of receptor activation as measured by tyrosine 594 and 604 phosphorylation.

**Figure 4.**
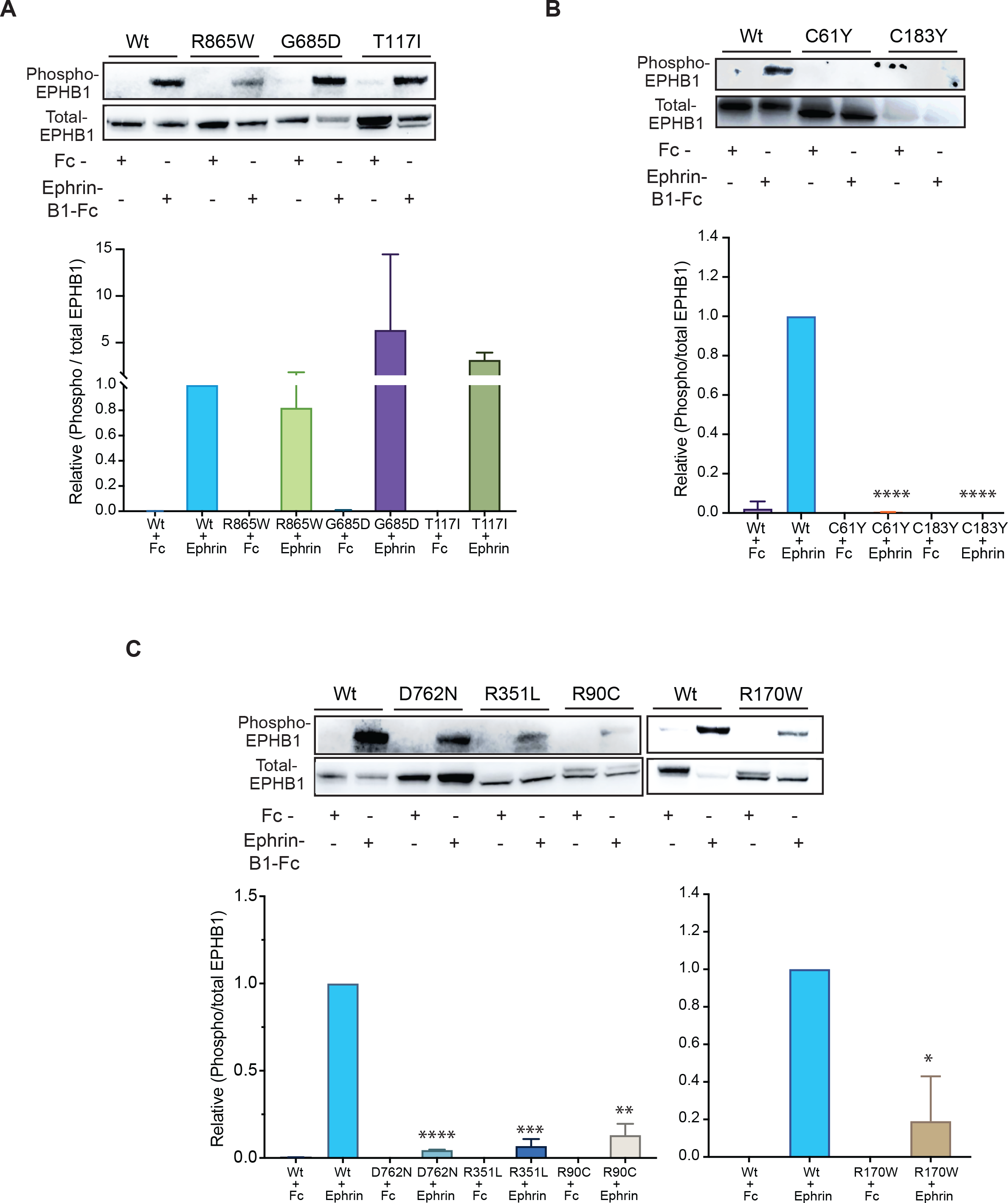
Phosphorylation of Y594/604 is reduced in *EPHB1* mutants with compromised or reduced compartmentalisation phenotypes. Representative immunoblot analyses on total protein from DLD-1 cell lines expressing EPHB1 mutants with (A) wild-type-like (R865W, G685D and T117I), (B) compromised (C61Y and C183Y) and (C) reduced compartmentalisation phenotype (D762N, R351L, R90C and R170W) stimulated with 0.5 μM EphrinB1-Fc ligand or negative control Fc fragment. Phosphorylation of EPHB1 Tyr594 and Tyr604 (upper panels) and total EPHB1 protein (middle panels) were detected with phospho-specific antibodies and FLAG-tag antibody, respectively. Phosphorylation was quantified as the ratio of band intensities of phosphorylated to total EPHB1 (lower panels) normalized to wild-type EPHB1 stimulated with ligand. Each experiment was repeated 3 times and the quantitation was based on results from all repetitions. Error bars, SD. The Mann-Whitney *U* test was used to calculate the *P*-values, * *p* < 0.05, ** *p* < 0.01, *** *p* < 0.001 and **** *p* < 0.0001.

### A mutation hotspot uncovered by 3D-proximity analysis

. Next, we hypothesised that our 3D-structure-based bioinformatic pipeline should be able to detect mutational hotspots in which both mutated amino acids in the pair show similar phenotypes. Among the identified mutants, *EPHB1* C61Y showed the most compromised phenotype as large clusters were almost absent (*p* < 0.0001; Figure 3A-B). In the TCGA dataset, *EPHB1* C61Y was found in a colon adenocarcinoma and, aligned in the same position of the protein consensus sequence, the corresponding *EPHA2* C70R mutant was found in a renal papillary cell carcinoma. The amino acid C61 was associated with the 3D-partner position C183, localised 122 amino acids C-terminally in *EPHB1* and 4.69 Å away in 3D-space (Figure 3C). Considering the same aligned amino acid position in the protein consensus sequence for other EPH receptors, we identified a urothelial cancer case with *EPHA2* C188Y mutation. To determine if the C61 partner position would have the same compromised phenotype, we engineered a cell line expressing *EPHB1* C183Y (NM_004441:c.G923A). The mutation was confirmed by Sanger sequencing at the transcript level and protein overexpression with immunoblotting (Supplementary Figures 2-3). Indeed, the C183Y mutant also had a strongly compromised compartmentalisation phenotype in the absence or presence of *EFNB1* ligand-expressing cells (*p* < 0.0001), and receptor phosphorylation after ligand stimulation was absent (Figure 4B). Taken together, the spatial proximity analysis uncovered hotspots not readily detectable by analyses of the linear EPH sequences.

### Proteome and phospho-proteome profile of ligand stimulated EPHB1 mutants

Next, we sought to identify alterations in signalling that could cause the different compartmentalisation phenotypes observed between EPHB1 mutations. We performed a combined proteome and phospho-proteome analysis of 1438 proteins in one mutant cell line representing each phenotype (i.e., C61Y (compromised), D762N and R351L (reduced with mutations in kinase and fibronectin-I domains, respectively), R743W (enhanced) and wildtype after stimulation with EphrinB1-Fc ligand. After ligand stimulation at 37°C for 30 min, 1-10 proteins and 7-64 differentially phosphorylated proteins per mutant were identified (Supplementary figure 6A-B). Ligand stimulation of EPHB1 wild-type cells resulted in 2 differentially expressed proteins (S10A8/9 and GLPB) and 1 phosphoprotein (TOP2A; Supplementary figure 7A-C). However, 0-7 differentially expressed proteins and 3-64 differentially phosphorylated proteins were identified in the four ligand-stimulated mutants as compared to stimulated wild-type control (Supplementary figure 8A-C; 9A-C; 10A-C and 11A-C). Whereas a single (ARHG2) or no differential protein was identified in ligand-stimulated wild-type vs EPHB1 D762N/R351L comparisons (Supplementary figure 9A and 10A), the number of differential phospho-proteins were comparatively higher than proteins in all comparisons (Supplementary figure 8B-C; 9B-C; 10B-C and 11B-C). As expected, the ligand-stimulated Wt vs mutant EPHB1 comparisons revealed fewer regulated proteins than the corresponding comparisons without stimulation (Supplementary figure 12A-D, *left panel*). Conversely, more differentially phosphorylated proteins were detected after ligand stimulation than without stimulation (Supplementary Figure 12E-H, *left panel*).

Furthermore, more mutant specific differential phospho-proteins than proteins were observed after ligand stimulation (Supplementary figure 12A-D and 12E-H). Next, we hypothesized that the proteins/phosphorylated proteins in the overlap regions of each of the Venn-analysis for ligand-stimulated ligand stimulated mutants compared to wildtype could represent proteins/phosphorylated proteins specifically regulated by each mutant. We found 4 (SELE, YEATS2, PRSS3 and ALB; *P*=3,22E-07, Hypergeometric distribution), 1 (MKI67) and 7 proteins (MKI67, CDKN3, MAPT, GPX1, CCNB1, CD72 and MUC17) in C61Y/R351L/R743W mutants, respectively (Supplementary figure 12A-D). On the contrary, 14 (ZBTB22, PIK3C2B, INSL4, EPHB1, MED27, SCGB1A1, TGFBR3, FGA, BIRC3, KLK3, RGMB, PRDX2, WIF1 and PIP5K1C; *P*=5,85E-23), 4 (PIK3C2B, RPL7 and CD53 ; *P*=8,47E-07), 2 (PIK3C2B and RPL7; *P*=4,19E-05) and 1 (RPL7; *P*=1,54E-02) differentially phosphorylated proteins were identified in ligand-stimulated C61Y/D762N/R351L/R743W mutants (Supplementary Figure 12E-H). We next asked whether phosphorylation of specific signalling proteins can explain the difference in compartmentalisation between mutants with compromised or reduced and WT or enhanced phenotype. Interestingly, PIK3C2B was more phosphorylated in all three mutants with compromised or reduced compartmentalisation (Supplementary figure 6-10A-C and 11E-H). Taken together, ligand stimulation of the different mutants elicited a higher degree of differential protein phosphorylation than differential proteins level, potentially because of the relatively short time between stimulation and cell lysis.

### Common and differential pathway enrichment after ligand stimulation

Next, we performed unbiased STRING based pathway analysis (https://string-db.org) (21) on both the differential phosphorylated proteins and on total proteins. In stimulated C61Y/D762N/R351L, PI3K-Akt (22) and immune-related pathways (e.g. interleukin, JAK-STAT, Toll-like receptor signaling and PD-1/PD-L1 pathways) (23) were commonly enriched (Supplementary figure 8-10D), MAPK and RAF/MAP kinase pathways in both C61Y/D762N (Supplementary figure 8-9D), HIF-1 pathways in R351L (Supplementary figure 10D), p53 signaling in C61Y (Supplementary figure 8D) and mTORC1, axon-guidance as well as Ribosomal pathways in R743W (Supplementary figure 11D) were differentially enriched as compared to stimulated Wt.

Next, we sought to understand how the mutant-specific phospho-proteome and proteome caused the different compartmentalisation phenotypes. In the ligand-stimulated C61Y mutant, the NF-kappa B and TNF pathways were enriched, both of which have been reported to cross-talk with Eph-ephrin signaling (Figure 5A) (24,25). Similarly, other pathways reported to cross-talk with Eph-ephrin signaling such as MAPK (26), gastric (27), colorectal cancer (13,28) and Ras-related pathways (29) were enriched in ligand-stimulated D762N kinase mutant with reduced compartmentalisation phenotype (Figure 5A). The fibronectin domain mutant R351L, with reduced compartmentalisation phenotype, had enrichment of HIF-1 (30), VEGF (31), and EGF/EGFR (32) (Figure 5A). Altogether, these pathways may contribute to the differential compartmentalisation phenotypes, but no pathways emerged as specific to any of the ligand-stimulated mutants (Figure 5B).

**Figure 5.**
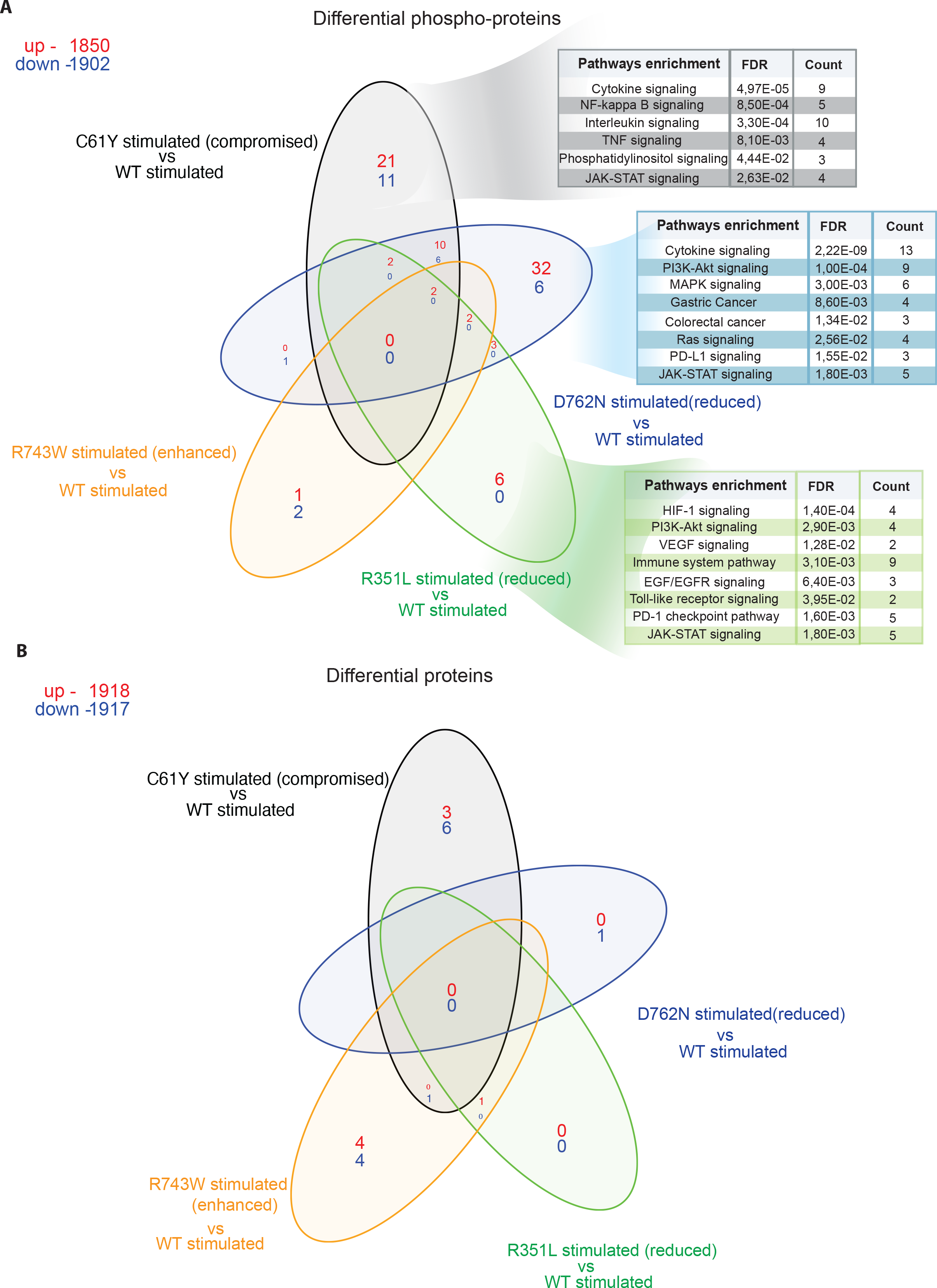
Enrichment of phosphatidylinositol and PI3K-Akt pathways in the phospho-proteomes of ligand stimulated *EPHB1* mutants with reduced or compromised compartmentalisation phenotypes. One mutant representing each category of compartmentalisation phenotype, C61Y (compromised), D762N (reduced), R351L (wild-type-like) and R743W (enhanced) along with wild-type EPHB1 were subjected to Ephrin-B1-Fc ligand stimulation for 30 min followed by array-based differential phospho-proteome (A) or total proteome (B) analysis. The Fc-stimulated wild-type EPHB1 served as negative control. Protein and phosphorylation levels of 1,438 different proteins were analysed using the Sciomics platform using 1,929 antibodies. Up- and down-regulated phospho-proteins and total proteins in red and blue, respectively. The enriched pathway from the STRING database (https://string-db.org) were shown on the differential phosphorylated proteins of EphrinB1-Fc ligand stimulated EPHB1 (C61Y/D762N/R351L) mutants.

## Discussion

A major challenge in understanding EPH receptor mutations observed in cancers is the combinatorial nature and bidirectional complexity of ephrin signalling. EPH receptor and EFN ligand-expressing cells interact with each other by repulsion to position themselves in the colonic crypts in a specific pattern along the crypt-villus axis (33). EPH receptors have previously been associated with metastatic disease development due to their role in tumour growth, invasiveness, angiogenesis and metastasis *in vivo* (19). While the role of specific EPH receptor mutations has been functionally studied in certain tumour types, like *EPHA3* in lung cancer, the role of *EPHB1* somatic mutations had not been studied to date (6,7). Due to the intermediate-to-low mutation frequencies of EPH receptors in cancer, we devised a strategy incorporating pan-cancer pan-EPH mutations and 3D-structure analyses of EPH receptors to maximise detection of mutational hotspots and clusters. After analysing pan-cancer and pan-EPH somatic mutational data from TCGA, we identified 8 mutations in 3D-cluster pairs and 7 recurrent mutated positions in *EPHB1* for further functional studies (Table 1). Large scale validation of EPH receptor mutations is challenging due to the elaborate nature of the established functional assays. The *EPHB2-EFNB1* interaction was shown to suppress colorectal cancer progression by compartmentalising tumour cells, which was substantiated in a mouse model (14). We recently demonstrated that *EPHB1*-*EFNB1* interactions can be studied in DLD-1 CRC cells using an *in vitro* compartmentalisation assay (5). Therefore, we selected compromised compartmentalisation as a semi-scalable read-out to identify *EPHB1* mutations with functional impact.

The N-terminal ligand-binding domain of *EPHB1* is responsible for the initiation of signal transduction after EFN ligand binds to this domain. Non-synonymous mutations can disrupt this function, which can abrogate the cellular repulsion caused by EPH-EFN interaction (34). This may explain the reduced compartmentalisation phenotype of the R90C, R170W, R351L and D762N mutants. Ligand stimulation experiments revealed significant reduction of R90C, R170W, R351L and D762N receptor phosphorylation level at Tyr592/604 compared to Wt, suggesting a reduced downstream signal transduction. Moreover, the C61Y mutation and its 3D-partner C183Y form a cysteine bridge in the ligand binding domain. The abrogation of this bond, which is of critical importance for domain structure, strongly compromised compartmentalisation with almost no large clusters observed in co-cultures with *EFNB1* expressing cells. This was accompanied by an absence of ligand-induced receptor Tyr592 and Tyr604 phosphorylation. However, an overall phosphorylation assay using a panel of 1929 antibodies spanning 1428 proteins as well as covering all of the serine, threonine and tyrosine phosphorylation positions identified 32 differential phosphorylated proteins after ligand stimulation, belonging to cancer relevant pathways such as cytokine, PI3K-Akt, MAPK, NF-Kappa ², Ras and JAK-STAT pathways.

The fibronectin domain of EPH receptors is responsible for the interactions with several other transmembrane proteins, such as integrins (35,36), and plays important roles in metastasis and invasion (37). Non-synonymous mutations in this domain can disrupt its function to interact with other proteins, and we previously demonstrated that *EPHB1* R351W significantly reduced compartmentalisation (5). Here, the R351L substitution at the same position also showed a reduced compartmentalisation phenotype as well as reduced ligand induced receptor Tyr592 and Tyr604 phosphorylation. Furthermore, the overall phosphorylation assay identified 6 upregulated phosphorylated proteins in cancer related pathways such as HIF-1, PI3K-Akt, EGF/EGFR, toll-like receptor, and JAK-STAT signalling.

Downstream signalling from *EPHB1* is transduced by phosphorylation of tyrosine residues in the kinase domain upon binding to EFNB ligands (38,39). The D762N mutant in the EPHB1 kinase domain had a reduced compartmentalisation phenotype in line with our previous study (5) and significant reduction of Tyr592 and Tyr604 phosphorylation level after ligand stimulation. Phospho-proteomics identified 38 differential phosphorylated proteins in the MAPK, Ras, Colorectal and Gastric cancer pathways.

In contrast, we observed enhanced compartmentalisation of the R743W, R748S and G821R kinase domain mutants along with reduced or absent receptor Tyr592 and Tyr604 phosphorylation, suggesting that a phosphorylation independent mechanism underlies enhanced compartmentalisation phenotypes. No altered pathways were identified in the phosphoproteome analysis. However, combined STRING analysis on native and phosphorylated proteins from ligand stimulated R743W showed enrichment of ribosomal, mTORC1 and axon-guidance pathways. Further research is warranted to find the mechanisms behind the enhanced compartmentalisation phenotype. Notably, PI3K-Akt pathways have previously been implicated in Eph-ephrin (22) and EPHB1 (5) signalling. Here, increased phosphorylation of PIK3C2B upon ligand stimulation characterized Eph receptor mutants with decreased compartmentalisation. In A-431 cells, a Grb2-PI3KC2B-Eps8-Abi1-Sos complex interacts with the EGF receptor and influences Rac activity, epithelial adherence junctions, and membrane ruffling (40). In the A-431 system, overexpression of PIK3C2B led to more compact colonies and in HEK293 cells to increased cell migration and altered actin reorganization on cell adhesion (41). A significant association between PIK3C2B and familial, early-onset prostate cancer has been observed (42). The PIK3C2B is therefore a plausible link between Eph receptor signalling and the clustering phenotype and is a strong candidate for future functional studies.

In conclusion, by accessing pan-cancer pan-EPH mutational data we were able to select 15 *EPHB1* mutants from which 7 mutants lacked impact, 2 enhanced, and 6 reduced or strongly compromised cell compartmentalisation. Whereas the 3D-protein structure-based bioinformatics analysis identified 63% (5 out of 8 selected mutants) *EPHB1* mutants with compartmentalisation phenotypes, the 2D-analysis identified 43% (3 out of 7 selected mutants), demonstrating the utility for 3D-protein structure-based mutation analysis in characterization of putative cancer genes. Further functional studies are warranted to establish mechanistic links between the compartmentalisation phenotype and metastatic disease development. This is, to date, the first study of pan-cancer *EPHB1* receptor mutations by an integrative approach involving 3D-protein structure-based bioinformatics analysis followed by *in vitro* validation, a robust way to identify cancer-causing mutations.

## Supplementary data statement

Supplementary Data are available at NAR online.

## Acknowledgement

The results shown here are in part based upon data generated by the TCGA Research Network: https://www.cancer.gov/tcga.

## Biosafety declaration

The Swedish work environment authority approved the work with genetically modified and replication deficient lentiviral particles (Arbetsmiljöverket ID 202100-2932 v72). All the experiments with GMO lentiviral particles were conducted under Biosafety Level 2.

