## Supplementary figures and tables for "Recurring *EPHB1* mutations in human cancers alter receptor signalling and compartmentalisation of colorectal cancer cells"

Running title: Phenotypes of *EPHB1* mutations in solid tumours

**Supplementary Figures:**

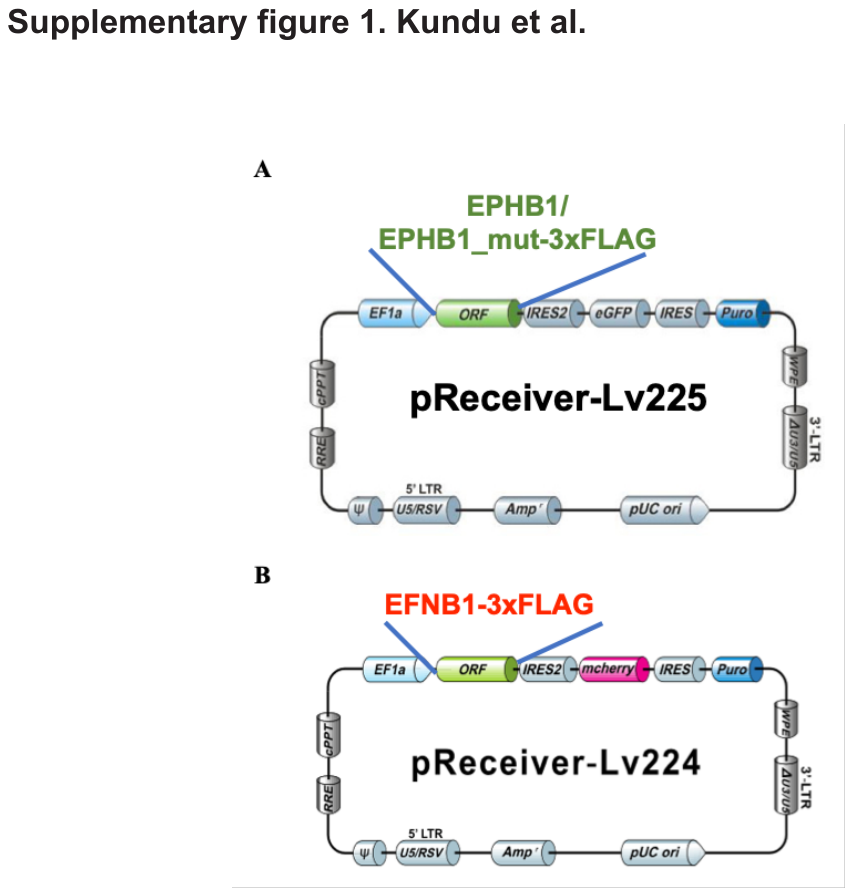

**Supplementary figure 1. Construct designs.** (A) Wild-type *EPHB1* and its mutants, and (B) wild-type *EFNB1* with a C-terminal 3×FLAG tag were cloned downstream of the EF1α-constitutive promoter in pReceiver-Lv225 and pReceiver-Lv224, respectively. EF1α, Human elongation factor-1 alpha mammalian constitutive promoter; ORF, open reading frame where different versions of *EPHB1* and *EFNB1* cDNAs were cloned; IRES and IRES2, internal ribosomal entry sites; eGFP, enhanced green fluorescent protein; mCherry, modified red fluorescent protein; Puro, puromycin-N-acetyltransferase gene; pUC ori, pUC replication of origin; Amp, beta-lactamase gene; cPPT, central polypurine tract; PRE, post-transcriptional regulatory element; WPE, Woodchuck hepatitis virus post-transcriptional regulatory element; 5’ and 3’LTR, 5’ and 3’ long terminal repeats.

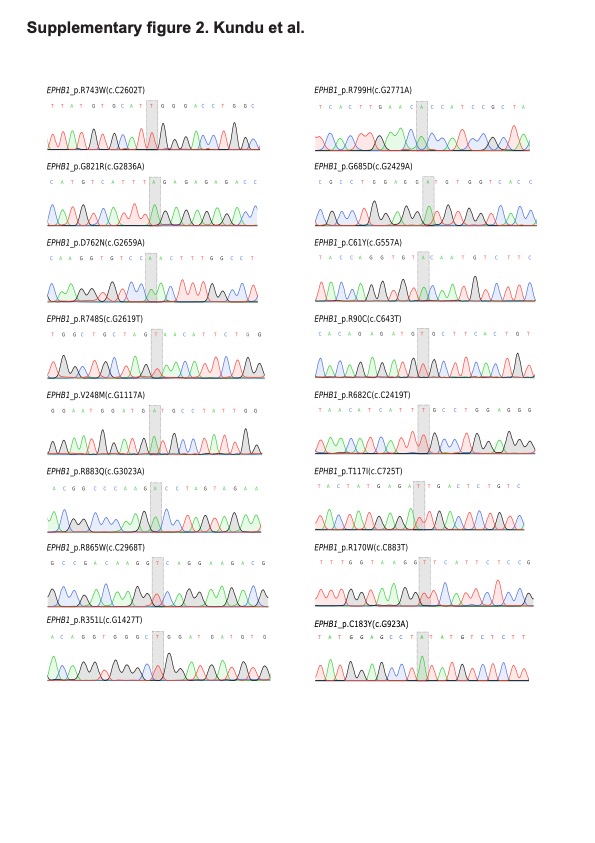

**Supplementary figure 2. Confirmation of *EPHB1* mutants at the transcript level by Sanger sequencing.** The mutation of interest is highlighted with a grey bar in each panel.

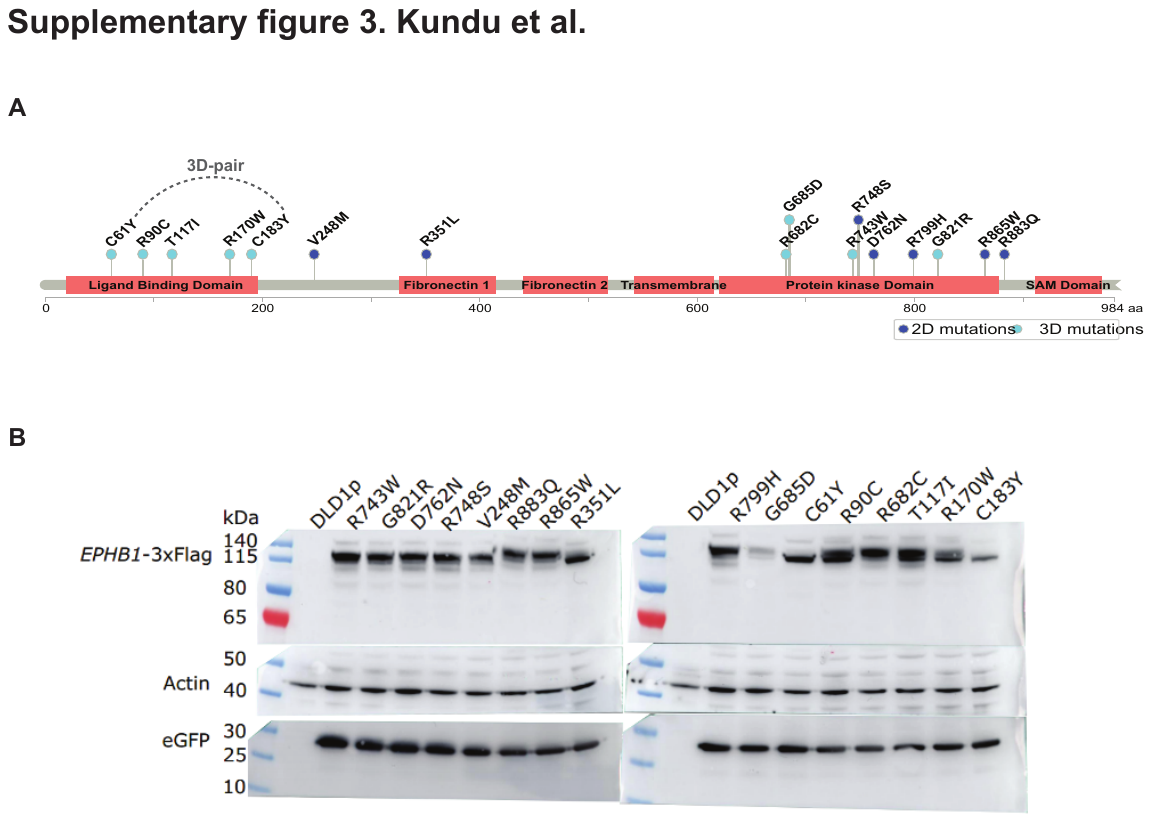

**Supplementary figure 3. Immunoblot analysis of ectopically expressed EPHB1 mutant proteins in DLD-1 CRC cells.** (A) All engineered EPHB1 mutations are depicted on the full-length (984 amino acids) EPHB1 protein. Mutants selected from 2D and 3D-structure based bioinformatic analysis in dark blue and light blue, respectively. The 3D-pairing between C61Y and C183Y is shown with a dotted line. (B) Expression of wild-type (positive control) and mutant EPHB1 proteins was detected with FLAG antibody (upper panel) with actin as a loading control (middle panel) and eGFP expression level was shown in each sample (lower panel). Parental DLD-1 cells (DLD1p) were used as a negative control for EPHB1 and eGFP expression.

**Supplementary figure 4. *EPHB1* receptor mutations T117I, G685D and R865W have wild type like response to *EFNB1* in compartmentalisation of colorectal cancer cells.** (A) Upper panel, positions of the mutants in the full length EPHB1 protein. Lower panel, *in vitro* compartmentalisation assays by co-culturing DLD-1 cells expressing eGFP along with either wild-type EPHB1 or mutants T117I, G685D or R865W with DLD-1 cells expressing mCherry with or without EFNB1 ligand. Representative confocal images from 10 randomly chosen image fields for each condition. Arrows, examples of large (>50 cells), homogeneous GFP^+^ cell clusters indicative of cell sorting and compartmentalisation. (B) Quantitation of the compartmentalisation experiments. Cell distribution was quantified by counting the percentage of GFP^+^ cells forming clusters of different sizes. In co-cultures of EFNB1 ligand with EPHB1 or its T117I, G685D or R865W mutants, similar high percentage of GFP^+^ cells were distributed into large homogeneous clusters of >50 cells. The Mann-Whitney *U* test was used to calculate the difference between each mutant with or without EFNB1 against the positive control wild-type EPHB1-type with respect to the formation of large clusters (>50 cells). This experiment was performed at least twice and imaged with five random fields from each experiment. Here, * *p* < 0.05, ** *p* < 0.01, *** *p* < 0.001 and **** *p* < 0.0001. Wt, wild-type.

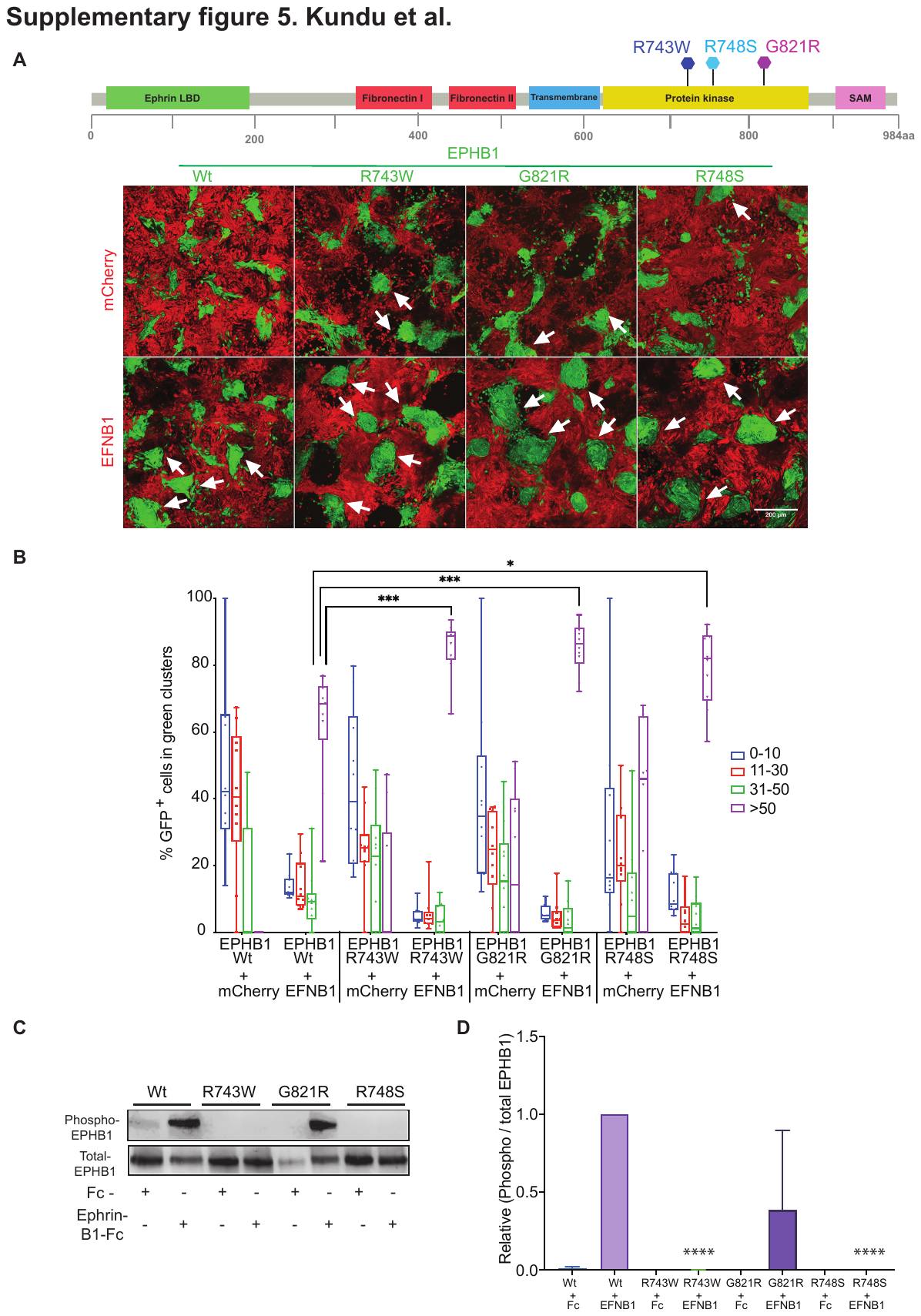

**Supplementary figure 5. The *EPHB1* enhanced compartmentalisation mutants R743W, G821R and R748S do not display increased Y594/604 phosphorylation in presence of *EFNB1*.** (A) Upper panel, positions of the mutants in the full length EPHB1 protein. Lower panel, *in vitro* compartmentalisation assays by co-culturing DLD-1 cells expressing eGFP along with either EPHB1 wild-type or R743W, G821R or R748S mutants with DLD-1 cells expressing mCherry with or without EFNB1 ligand. Representative confocal images from 10 randomly chosen image fields for each condition. Arrows, examples of large, homogeneous GFP^+^ cell clusters (>50 cells) indicative of cell sorting and compartmentalisation. (B) Quantitative results of the compartmentalisation experiments. Cell distribution was quantified by counting the percentage of GFP^+^ cells forming clusters of different sizes. (C) The levels of Y594/604 phosphorylation (upper panel) and total EPHB1 (lower panel) after Ephrin-B1-Fc ligand stimulations were analyzed by immunoblotting using antibodies to phospho-Y594/604 and total EPHB1 protein. (D) The phosphorylation level of wild-type and mutant EPHB1 was computed as the ratio of band intensities of phosphorylated to total EPHB1 normalised to wild-type from three independent experiments. Error bars, SD. The unpaired t-test was used to calculate the *P*-values for the normalized level of phosphorylation relative to wild-type + EphrinB1-Fc sample, * *p* < 0.05, ** *p* < 0.01, *** *p* < 0.001 and **** *p* < 0.0001. Wt, wild-type.

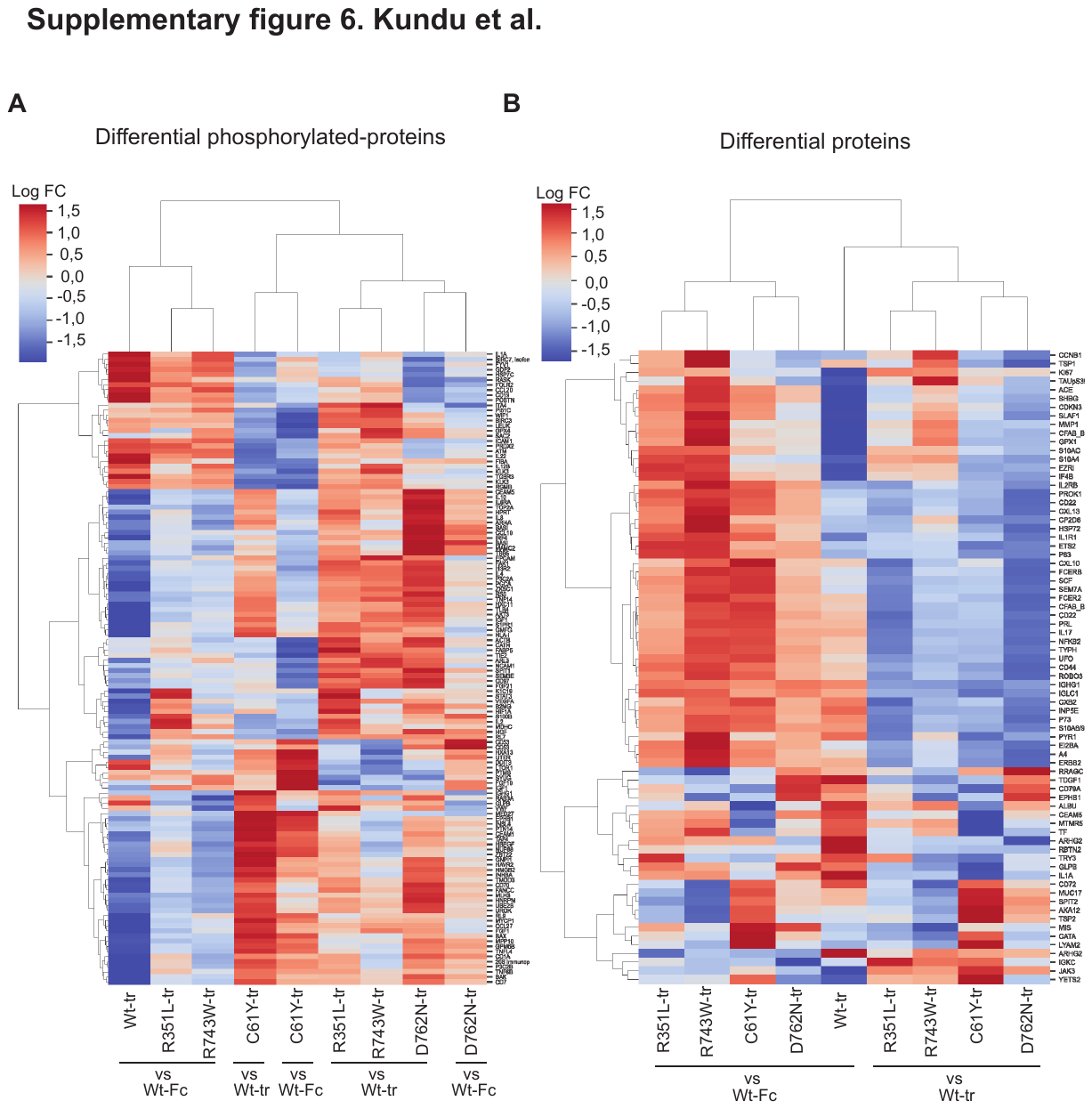

**Supplementary figure 6. Differential expression and phosphorylation of proteins after EphrinB1-Fc ligand stimulations.** An antibody-array-based method (Sciomics) was used with three replicates for each sample stimulated with 0.5 µM of EphrinB1-Fc ligand for 30 min of incubation. Heatmap represented differentially phosphorylated (A) and total (B) proteins with |logFC| > 1.0, adjusted *p* value < 0.05 in Ephrin-B1-Fc ligand stimulated wild-type EPHB1 and R351L, R743W, C61Y and D762N mutants in DLD-1 cells. Samples and genes clustered by Euclidian distance and Ward variance minimization with row-wise scaling.

**Supplementary figure 7. Total protein and phospho-protein profiles of EphrinB1-Fc ligand stimulated wild-type EPHB1 expressing DLD-1 cells.** An antibody-array-based method (Sciomics) was used with three replicates for each sample stimulated with 0.5 μM of EphrinB1-Fc ligand for 30 min for differential proteome (A), phospho-proteome (B), and joint (C) analysis. Differences in abundance between EphrinB1-Fc ligand and control Fc fragment stimulated EPHB1 expressing cells as log-fold change (logFC) and corresponding *p* values adjusted for multiple testing. Red line, adjusted *p* value significance level of 0.05; vertical lines, logFC cutoffs of ± 0.5. A positive and negative logFC indicates higher abundance or phosphorylation in ligand and Fc-stimulated wild-type EPHB1, respectively. Blue symbols, proteins differentially expressed or phosphorylated, defined as |logFC| > 0.5 and adjusted *p* value < 0.05. Green symbols, noteworthy proteins including non-significant proteins (adjusted *p* value < 0.9) with |logFC| > 1, and significant proteins with |logFC| ± 0.25. WT, wild-type.

**Supplementary figure 8. Differential protein expression and phosphorylation after ligand stimulation of C61Y versus wild-type EPHB1 DLD-1 cells.** An antibody-array-based method (Sciomics) was used with three replicates for each sample stimulated with 0.5 μM of EphrinB1-Fc ligand for 30 min for differential proteome (A), phospho-proteome (B), and joint (C) analysis. Differences in abundance between C61Y-stimulated samples and WT-stimulated samples as log-fold changes (logFC) and their corresponding *p* values (adjusted for multiple testing). Red line, adjusted *p* value significance level of 0.05; vertical lines, logFC cut-offs of ± 0.5. A positive and negative logFC indicates higher abundance or phosphorylation in ligand-stimulated C61Y mutant and wild-type EPHB1, respectively. Blue symbols, proteins differentially expressed or phosphorylated, defined as |logFC| > 0.5 and adjusted *p* value < 0.05. Green symbols, noteworthy proteins including non-significant proteins (adjusted *p* value < 0.9) with |logFC| > 1, and significant proteins with |logFC| ± 0.25. (D) Pathway analysis was performed with only the differential proteins/phosphorylated-proteins (blue) using STRING analysis (<https://string-db.org>) setting criteria full STRING network; Confidence network edges and high confidence (7.00). WT, wild-type.

**Supplementary figure 9. Differential protein expression and phosphorylation after ligand stimulation of D762N versus wild-type EPHB1 DLD-1 cells.** An antibody-array-based method (Sciomics) was used with three replicates for each sample stimulated with 0.5 μM of EphrinB1-Fc ligand for 30 min for differential proteome (A), phospho-proteome (B), and joint (C) analysis. Differences in abundance between D762N-stimulated samples and WT-stimulated samples as log-fold changes (logFC) and their corresponding *p* values (adjusted for multiple testing).Red line, adjusted *p* value significance level of 0.05; vertical lines, logFC cutoffs of ± 0.5. A positive and negative logFC indicates higher abundance or phosphorylation in ligand-stimulated D762N mutant and wild-type EPHB1, respectively. Blue symbols, proteins differentially expressed or phosphorylated, defined as |logFC| > 0.5 and adjusted *p* value < 0.05. Green symbols, noteworthy proteins including non-significant proteins (adjusted *p* value < 0.9) with |logFC| > 1, and significant proteins with |logFC| ± 0.25. (D) Pathway analysis was performed with only the differential proteins/phosphorylated-proteins (blue) using STRING analysis (<https://string-db.org>) setting criteria like, full STRING network; Confidence network edges and high confidence (7.00). WT, wild-type.

**Supplementary figure 10. Differential protein expression and phosphorylation after ligand stimulation of R351L versus wild-type EPHB1 DLD-1 cells.** An antibody-array-based method (Sciomics) was used with three replicates for each sample stimulated with 0.5 μM of EphrinB1-Fc ligand for 30 min for differential proteome (A), phospho-proteome (B), and joint (C) analysis. Differences in abundance between R351L-stimulated samples and WT-stimulated samples as log-fold changes (logFC) and their corresponding p values (adj. for multiple testing). Red line, adjusted p value significance level of 0.05; vertical lines, logFC cutoffs of ± 0.5. A positive and negative logFC indicates higher abundance or phosphorylation in ligand-stimulated R351L mutant and wild-type EPHB1, respectively. Blue symbols, proteins differentially expressed or phosphorylated, defined as |logFC| > 0.5 and adjusted p value < 0.05. Green symbols, noteworthy proteins including non-significant proteins (adj. p value < 0.9) with |logFC| > 1, and significant proteins with |logFC| ± 0.25. (D) Pathway analysis was performed with only the differential proteins/phosphorylated-proteins (blue) using STRING analysis (<https://string-db.org>) setting criteria like, full STRING network; Confidence network edges and high confidence (7.00).

**Supplementary figure 11. Differential protein expression and phosphorylation after ligand stimulation of R743W versus wild-type EPHB1 DLD-1 cells.** An antibody-array-based method (Sciomics) was used with three replicates for each sample stimulated with 0.5 μM of EphrinB1-Fc ligand for 30 min for differential proteome (A), phospho-proteome (B), and joint (C) analysis. Differences in abundance between R743W-stimulated samples and WT-stimulated samples as log-fold changes (logFC) and their corresponding *p* values (adjusted for multiple testing). Red line, adjusted *p* value significance level of 0.05; vertical lines, logFC cutoffs of ± 0.5. A positive and negative logFC indicates higher abundance or phosphorylation in ligand-stimulated R743W mutant and wild-type EPHB1, respectively. Blue symbols, proteins differentially expressed or phosphorylated, defined as |logFC| > 0.5 and adjusted *p* value < 0.05. Green symbols, noteworthy proteins including non-significant proteins (adjusted *p* value < 0.9) with |logFC| > 1, and significant proteins with |logFC| ± 0.25. (D) Pathway analysis was performed with only the differential proteins/phosphorylated-proteins (blue) using STRING analysis (<https://string-db.org>) setting criteria like, full STRING network; Confidence network edges and high confidence (7.00). WT, wild-type.

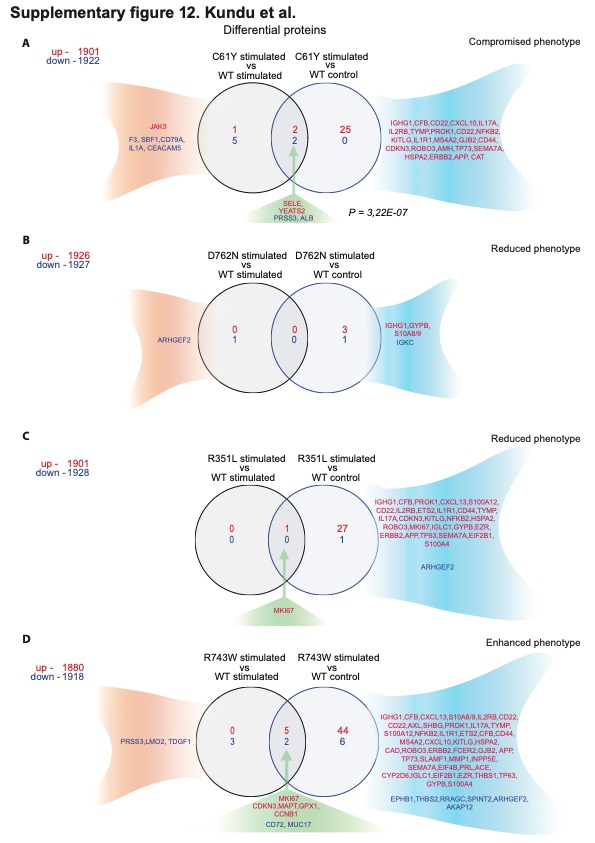

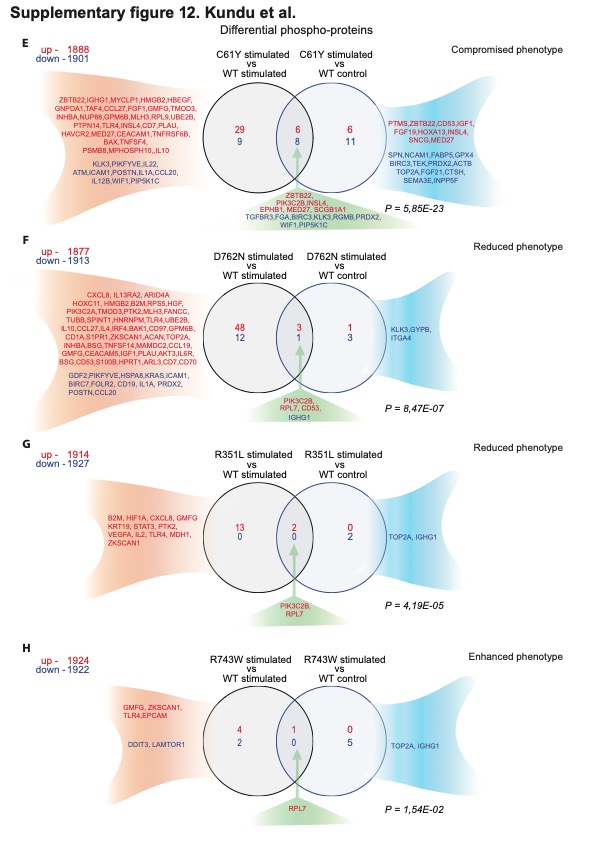

**Supplementary figure 12. Greater differential protein phosphorylation than differential protein expression in ligand stimulated mutants versus wild-type EPH receptor.** After stimulating DLD1 cells ectopically expressing EPHB1 wild-type and C61Y/D762N/R351L/R743W mutants with EphrinB1-Fc ligand or Fc (control), they were subjected to array-based total proteome and Phosphoproteome analysis (Sciomics) to identify differential proteins (A-D) and phospho-proteins (E-H). Up, upregulation (red); Down, downregulation (blue). The hypergeometric distribution was used to calculate the *p* values. WT, wild-type.

**Supplementary Tables:**

| Supplementary Table 1. Calculation of customised average functional impact. | | |
| --- | --- | --- |
|  | **Score +1** | **Score +0.5** |
| Functional Prediction Scores: |  |  |
| SIFT | “damaging (D)” | - |
| Polyphen2_HDIV | “probably damaging (D)” | “possibly damaging (P)” |
| Polyphen2_HVAR | “probably damaging (D)” | “possibly damaging (P)” |
| LRT | “deleterious (D)” | - |
| MutationTaster | “disease causing automatic (A)” | “disease causing (D)” |
| MutationAssessor | “predicted functional high (H)” | “predicted functional medium (M)” |
| FATHMM | “damaging (D)” | - |
| PROVEAN | “deleterious (D)” | - |
| Fathmm_MKL | “deleterious (D)” | - |
| Ensemble Scores: |  |  |
| MetaSVM | “deleterious (D)” | - |
| MetaLR | “deleterious (D)” | - |
| M-CAP | “deleterious (D)” | - |
| CADD | >= 30 | **-** |

| Supplementary Table 2. Identifiers of EPH receptors protein and Protein Data Bank 3D-structural entries. | | |
| --- | --- | --- |
| Ephrin Receptor | **UniProt** | **PDB Files Entries** |
| *EPHA1* | P21709 | 2K1K 2K1L 3HIL 3KKA |
| *EPHA2* | P29317 | 1MQB 2E8N 2K9Y 2KSO 2X10 2X11 3C8X 3CZU 3FL7 3HEI 3HPN 3KKA 3MBW 3MX0 3SKJ 4P2K 4PDO 4TRL 5EK7 5I9U 5I9V 5I9W 5I9X 5I9Y 5I9Z 5IA0 5IA1 5IA2 5IA3 5IA4 5IA5 5NJZ 5NK0 5NK1 5NK2 5NK3 5NK4 5NK5 5NK6 5NK7 5NK8 5NK9 5NKA 5NKB 5NKC 5NKD 5NKE 5NKF 5NKG 5NKH 5NKI |
| *EPHA3* | P29320 | 2GSF 2QO2 2QO7 2QO9 2QOB 2QOC 2QOD 2QOF 2QOI 2QOK 2QOL 2QON 2QOO 2QOQ 3DZQ 3FXX 3FY2 4G2F 4GK2 4GK3 4GK4 4L0P 4P4C 4P5Q 4P5Z 4TWN 4TWO |
| *EPHA4* | P54764 | 2LW8 2WO1 2WO2 2WO3 3CKH 3GXU 4BK4 4BK5 4BKA 4BKF 4M4P 4M4R 4W4Z 4W50 5JR2 |
| *EPHA5* | P54756 | 2R2P 4ET7 |
| *EPHA6* | Q9UF33 | No structures available |
| *EPHA7* | Q15375 | 2REI 3DKO 3H8M 3NRU |
| *EPHA8* | P29322 | 1UCV 1X5L 3KUL |
| *EPHA10* | Q5JZY3 | No structures available |
| *EPHB1* | P54762 | 2DJS 2EAO 3ZFX 5MJA 5MJB |
| *EPHB2* | P29323 | 1B4F 1F0M 2QBX 3ZFM |
| *EPHB3* | P54753 | 3P1I 3ZFY 5L6O 5L6P |
| *EPHB4* | P54760 | 2BBA 2E7H 2HLE 2QKQ 2VWU 2VWV 2VWW 2VWX 2VWY 2VWZ 2VX0 2VX1 2X9F 2XVD 2YN8 3ZEW 4AW5 4BB4 |
| *EPHB6* | O15197 | No structures available |

| Supplementary Table 3. PCR primers for Sanger sequencing for *EPHB1* mutant detection. | | | | |
| --- | --- | --- | --- | --- |
| Primer ID | **Sequence (5’ -> 3’)** | **Tmº** | **Protein Start Position** | **Product Size (bp)** |
| #1-Forward | AGTGGCTGCGATGGAAGAAAC | 67.5 | 42 | 636 |
| #1-Reverse | CCCGAGCAATCACCAGAGATG | 69.1 | 677 |  |
| #2-Forward | CTTCAAAAAGTGTCCCAGCATTG | 66.5 | 586 | 644 |
| #2-Reverse | GTCAAAGGTGTAGGGGGTGTG | 65.4 | 1,229 |  |
| #3-Forward | GCAGGGGAGTTTGGAGAAGTG | 67.1 | 1,893 | 943 |
| #3-Reverse | CAGTGAGGAAGCTGTCCCTG | 65.1 | 2,835 |  |
| #4-Forward | GAGAATGGTGCATTGGATTCTTTC | 66.5 | 2,119 | 500 |
| #4-Reverse | CCTTCTGCCAACAGTCCAGCATG | 71.9 | 2,618 |  |

| Supplementary Table 4. Description of the 33 tumour types used by total amount of patients and EPH receptor mutations. | | | |
| --- | --- | --- | --- |
| Abbreviation | **Tumour Type Name** | **Number of Patients** | **Number of Mutations** |
| ACC | Adrenocortical carcinoma | 83 | 318 |
| BLCA | Bladder urothelial carcinoma | 389 | 1,795 |
| BRCA | Breast invasive carcinoma | 994 | 5,993 |
| CESC | Cervical squamous cell carcinoma and endocervical adenocarcinoma | 298 | 1,654 |
| CHOL | Cholangiocarcinoma | 50 | 267 |
| COAD | Colon adenocarcinoma | 419 | 8,624 |
| DLBC | Lymphoid neoplasm diffuse large B-cell lymphoma | 47 | 761 |
| ESCA | Esophageal carcinoma | 174 | 766 |
| GBM | Glioblastoma multiforme | 388 | 3,440 |
| HNSC | Head and neck squamous cell carcinoma | 490 | 2,565 |
| KICH | Kidney chromophobe | 64 | 230 |
| KIRC | Kidney renal clear cell carcinoma | 334 | 2,936 |
| KIRP | Kidney renal papillary cell carcinoma | 276 | 1,589 |
| LAML | Acute myeloid leukemia | 146 | 2,004 |
| LGG | Brain lower grade glioma | 443 | 1,447 |
| LIHC | Liver hepatocellular carcinoma | 368 | 3,103 |
| LUAD | Lung adenocarcinoma | 554 | 4,810 |
| LUSC | Lung squamous cell carcinoma | 485 | 4,291 |
| MESO | Mesothelioma | 81 | 348 |
| OV | Ovarian serous cystadenocarcinoma | 436 | 4,329 |
| PAAD | Pancreatic adenocarcinoma | 183 | 1,020 |
| PCPG | Pheochromocytoma and paraganglioma | 133 | 267 |
| PRAD | Prostate adenocarcinoma | 429 | 1,447 |
| READ | Rectum adenocarcinoma | 154 | 2,763 |
| SARC | Sarcoma | 245 | 1,007 |
| SKCM | Skin cutaneous melanoma | 464 | 7,149 |
| STAD | Stomach adenocarcinoma | 422 | 3,016 |
| TGCT | Testicular germ cell tumours | 146 | 739 |
| THCA | Thyroid carcinoma | 452 | 2,103 |
| THYM | Thymoma | 118 | 420 |
| UCEC | Uterine corpus endometrial carcinoma | 522 | 7,616 |
| UCS | Uterine carcinosarcoma | 53 | 221 |
| UVM | Uveal melanoma | 58 | 113 |

*, mutations selected for *in vitro* studies; ^1^distance between the amino acids in the pair < 5 Å; ^2^distance between the pair in the linear protein sequence > 50 amino acids; ^3^average functional impact score ≥ 6.

*Selected mutations for *in vitro* studies; ^1^distance between the pair less than 5 Å; ^2^distance between the pair more than 50 amino acids; ^3^average functional impact score more or equal than six.

| Supplementary Table 5. Alteration frequencies per Ephrin receptor according to tumour type and ratio of coding non-synonymous mutations per amino acid. | | | | | | | | | | | | | | | |
| --- | --- | --- | --- | --- | --- | --- | --- | --- | --- | --- | --- | --- | --- | --- | --- |
| Tumours | ***EPHA1*** | ***EPHA2*** | ***EPHA3*** | ***EPHA4*** | ***EPHA5*** | ***EPHA6*** | ***EPHA7*** | ***EPHA8*** | ***EPHA10*** | ***EPHB1*** | ***EPHB2*** | ***EPHB3*** | ***EPHB4*** | ***EPHB6*** | **Total** |
| SKCM | 6.0 | 5.6 | 10.8 | 8.0 | 3.7 | 15.3 | 13.4 | 5.2 | 5.2 | 5.8 | 9.3 | 6.5 | 3.2 | 8.6 | 51.1 |
| DLBC | 4.3 | 2.1 | 12.8 | 2.1 | 6.4 | 6.4 | 4.3 | 2.1 | 10.6 | 2.1 | 8.5 | 2.1 | 8.5 | 4.3 | 40.4 |
| LUAD | 1.3 | 1.8 | 7.4 | 3.2 | 11.0 | 7.0 | 6.3 | 2.2 | 2.3 | 6.3 | 2.3 | 3.2 | 1.6 | 7.4 | 38.4 |
| LUSC | 1.6 | 1.9 | 6.8 | 2.3 | 9.1 | 6.6 | 4.5 | 2.5 | 1.6 | 4.7 | 1.9 | 3.3 | 1.9 | 3.3 | 37.7 |
| STAD | 2.4 | 6.2 | 5.7 | 4.3 | 6.6 | 5.5 | 3.8 | 2.8 | 3.3 | 4.7 | 3.3 | 2.6 | 1.9 | 4.3 | 31.3 |
| UCEC | 4.6 | 5.6 | 8.0 | 8.8 | 9.8 | 8.0 | 6.1 | 4.0 | 4.4 | 10.9 | 5.0 | 3.6 | 4.2 | 4.6 | 31.0 |
| COAD | 1.2 | 3.1 | 6.9 | 3.6 | 6.9 | 5.7 | 4.5 | 2.9 | 3.1 | 6.0 | 3.8 | 2.1 | 1.7 | 2.9 | 30.8 |
| CRC | 1.0 | 2.4 | 5.8 | 3.3 | 6.8 | 5.1 | 4.2 | 2.8 | 2.6 | 5.6 | 2.8 | 1.7 | 1.9 | 2.8 | 27.9 |
| BLCA | 1.0 | 5.1 | 4.6 | 0.8 | 4.4 | 2.3 | 2.6 | 1.5 | 1.5 | 3.3 | 2.8 | 1.8 | 1.0 | 3.1 | 26.7 |
| HNSC | 1.4 | 4.9 | 2.9 | 1.2 | 4.3 | 2.2 | 3.7 | 1.2 | 1.4 | 2.2 | 0.8 | 1.2 | 1.4 | 1.4 | 22.4 |
| READ | 0.6 | 0.6 | 2.6 | 2.6 | 6.5 | 3.2 | 3.2 | 2.6 | 1.3 | 4.5 | 0.0 | 0.6 | 2.6 | 2.6 | 20.1 |
| ESCA | 0.0 | 2.9 | 1.7 | 1.7 | 4.0 | 2.3 | 1.1 | 2.3 | 1.1 | 1.7 | 2.9 | 0.6 | 0.6 | 1.7 | 20.1 |
| CESC | 0.7 | 4.0 | 1.3 | 1.3 | 1.7 | 1.7 | 1.0 | 2.3 | 1.3 | 1.3 | 2.0 | 1.0 | 2.0 | 1.7 | 17.4 |
| LIHC | 0.8 | 1.1 | 1.9 | 3.5 | 0.8 | 1.9 | 1.4 | 1.4 | 0.8 | 1.4 | 1.4 | 1.6 | 0.3 | 0.5 | 16.0 |
| GBM | 1.8 | 0.8 | 2.3 | 1.3 | 1.3 | 1.3 | 2.1 | 2.3 | 2.1 | 1.8 | 1.8 | 0.3 | 1.5 | 2.8 | 15.5 |
| OV | 1.1 | 1.4 | 1.6 | 0.7 | 1.4 | 1.4 | 1.6 | 0.7 | 0.9 | 1.4 | 1.4 | 0.9 | 0.9 | 2.3 | 14.4 |
| UCS | 1.9 | 0.0 | 1.9 | 1.9 | 5.7 | 0.0 | 0.0 | 0.0 | 0.0 | 3.8 | 3.8 | 0.0 | 0.0 | 0.0 | 11.3 |
| CHOL | 0.0 | 2.0 | 0.0 | 2.0 | 2.0 | 0.0 | 0.0 | 2.0 | 0.0 | 0.0 | 2.0 | 0.0 | 0.0 | 2.0 | 10.0 |
| MESO | 1.2 | 0.0 | 1.2 | 0.0 | 2.5 | 0.0 | 1.2 | 0.0 | 1.2 | 0.0 | 1.2 | 0.0 | 1.2 | 0.0 | 9.9 |
| BRCA | 0.7 | 0.5 | 0.7 | 1.2 | 1.3 | 1.0 | 1.9 | 0.1 | 0.3 | 1.8 | 0.5 | 1.1 | 0.8 | 0.9 | 9.9 |
| SARC | 0.4 | 0.8 | 0.4 | 0.8 | 2.0 | 0.8 | 1.6 | 1.2 | 0.4 | 0.8 | 0.4 | 0.0 | 0.4 | 0.4 | 9.4 |
| KIRP | 0.0 | 1.1 | 1.4 | 0.7 | 0.4 | 0.7 | 1.1 | 0.7 | 0.0 | 0.4 | 0.0 | 0.7 | 1.1 | 0.7 | 8.3 |
| ACC | 1.2 | 0.0 | 0.0 | 1.2 | 1.2 | 0.0 | 0.0 | 1.2 | 1.2 | 1.2 | 0.0 | 1.2 | 1.2 | 1.2 | 7.2 |
| LAML | 0.0 | 0.0 | 2.1 | 0.7 | 0.7 | 0.0 | 0.7 | 0.0 | 0.7 | 0.0 | 1.4 | 0.0 | 0.7 | 0.7 | 6.8 |
| KIRC | 0.6 | 0.3 | 0.3 | 0.3 | 1.5 | 0.6 | 0.3 | 0.3 | 0.3 | 0.3 | 0.0 | 0.3 | 0.6 | 0.6 | 6.3 |
| LGG | 0.5 | 0.5 | 1.4 | 1.1 | 0.7 | 0.0 | 0.9 | 0.9 | 0.7 | 0.0 | 0.5 | 0.5 | 0.5 | 0.5 | 5.4 |
| PAAD | 0.5 | 1.1 | 0.0 | 1.1 | 1.6 | 0.5 | 1.6 | 1.1 | 0.5 | 0.0 | 0.5 | 1.1 | 0.5 | 0.5 | 4.9 |
| TGCT | 0.0 | 0.0 | 0.7 | 0.0 | 0.0 | 1.4 | 0.0 | 0.0 | 1.4 | 0.0 | 0.0 | 0.0 | 2.1 | 0.0 | 4.8 |
| KICH | 1.6 | 0.0 | 1.6 | 0.0 | 0.0 | 0.0 | 0.0 | 0.0 | 1.6 | 1.6 | 0.0 | 0.0 | 0.0 | 0.0 | 4.7 |
| PRAD | 0.7 | 0.9 | 0.7 | 0.2 | 0.0 | 0.2 | 0.2 | 0.2 | 0.5 | 0.9 | 0.7 | 0.5 | 0.0 | 0.2 | 4.2 |
| PCPG | 0.0 | 0.0 | 0.8 | 0.0 | 0.0 | 0.8 | 0.0 | 0.0 | 0.0 | 0.0 | 0.0 | 0.0 | 0.0 | 2.3 | 3.8 |
| THCA | 0.4 | 0.0 | 0.0 | 0.0 | 0.2 | 0.4 | 0.0 | 0.4 | 0.7 | 0.7 | 0.2 | 0.0 | 0.2 | 0.2 | 3.3 |
| THYM | 0.0 | 0.0 | 0.0 | 0.0 | 0.0 | 0.0 | 0.8 | 0.0 | 0.0 | 0.0 | 0.0 | 1.7 | 0.0 | 0.0 | 2.5 |
| UVM | 0.0 | 0.0 | 0.0 | 0.0 | 1.7 | 0.0 | 0.0 | 0.0 | 0.0 | 0.0 | 0.0 | 0.0 | 0.0 | 0.0 | 1.7 |
| Mut/aa* | 0.15 | 0.25 | 0.40 | 0.26 | 0.39 | 0.38 | 0.35 | 0.18 | 0.17 | 0.33 | 0.21 | 0.17 | 0.15 | 0.26 | - |

* Ratio of coding non-synonymous mutations per amino acid protein length for each Ephrin receptor.

* Ratio of coding non-synonymous mutations per amino acid protein length for each Ephrin receptor.

*Selected mutations for *in vitro* studies; ^1^distance between the pair < 5 Å; ^2^distance between the pairs > 50 amino acids; ^3^average functional impact score ≥ 6.

| Supplementary Table 6. Ranked 3D-mutation in *EPHB1* sorted by average functional impact. | | | | | | | |
| --- | --- | --- | --- | --- | --- | --- | --- |
| ***EPHB1* Amino acid Position** | ***EPHB1***  **Mutations** | **Total EPH Mutations** | **3D Partner Position**  **(*EPHB1*)** | **3D Partner EPH** **Mutations** | **Pair Distance (Å)^1^** | **Pair Distance (Amino acids)^2^** | **Average Functional**  **Impact^3^** |
| **G685*** | 1 | 4 | I696 | 3 | 4.03 | 53 | 11.5 |
| **R743*** | 5 | 13 | F801 | 2 | 4.87 | 58 | 11.2 |
| **G821*** | 2 | 7 | R170 | 7 | 4.43 | 111 | 11 |
| V391 | 1 | 3 | L340 | 1 | 4.85 | 58 | 10.5 |
| L709 | 1 | 2 | F820 | 1 | 4.59 | 111 | 10.5 |
| **C61*** | 1 | 2 | C183 | 1 | 4.69 | 147 | 10 |
| **R90*** | 2 | 6 | S188 | 2 | 4.11 | 118 | 8 |
| M23 | 1 | 3 | V191 | 3 | 4.55 | 195 | 7 |
| **R682*** | 3 | 10 | E698 | 1 | 4.45 | 58 | 6.8 |
| **T117*** | 2 | 3 | G172 | 3 | 4.44 | 73 | 6.3 |
| **R170*** | 7 | 9 | E116 | 5 | 4.72 | 72 | 5.9 |

*Selected mutations for *in vitro* studies; ^1^distance between the pair < 5 Å; ^2^distance between the pairs > 50 amino acids; ^3^average functional impact score ≥ 6.

| Supplementary Table 7. Top 20 *EPHB1* mutations with no 3D-partner ranked by total EPH receptor mutations. | | | | |
| --- | --- | --- | --- | --- |
| ***EPHB1* Amino acid Position** | ***EPHB1***  **Mutations** | | **Total EPH Mutations** | **Average Functional**  **Impact^1^** |
| **V248*** | 1 | 26 | | 9 |
| **R883*** | 1 | 18 | | 8 |
| **R748*** | 2 | 14 | | 10 |
| **D762*** | 6 | 14 | | 11.9 |
| **R865*** | 3 | 13 | | 8.5 |
| D374 | 1 | 12 | | 6 |
| **R351*** | 2 | 11 | | 11.5 |
| **R799*** | 1 | 10 | | 10.5 |
| R304 | 3 | 10 | | 9.7 |
| R84 | 2 | 10 | | 9 |
| N290 | 8 | 10 | | 7.5 |
| D806 | 3 | 9 | | 12.5 |
| R767 | 1 | 9 | | 12 |
| P249 | 1 | 9 | | 10.5 |
| R663 | 3 | 9 | | 10 |
| D615 | 1 | 9 | | 9.5 |
| E131 | 1 | 9 | | 8 |
| E613 | 1 | 9 | | 7.5 |
| R953 | 2 | 9 | | 7.5 |
| R56 | 2 | 9 | | 7.25 |

*Selected mutations for *in vitro* studies; ^1^average functional impact score ≥ 6.
